# Cross-platform Hi-C meta-analysis identifies functional insulators that actively block enhancer-promoter interactions

**DOI:** 10.1101/2025.05.26.656183

**Authors:** Jian Cui, Wanying Xu, Xiuyuan Lang, Shanshan Zhang, Leina Lu, Xiaoxiao Liu, Yan Li, Fulai Jin

## Abstract

CTCF forms the boundaries of topologically associating domains (TADs) but their insulating function in transcription is controversial. Here we report that functional insulators (FINs) shall be determined not only by their ability to form static loops or boundaries, but also by the dynamic context if their flanking sequences form new loops upon boundary loss. We performed a meta-analysis of nine independent Hi-C and micro-C datasets upon acute CTCF- or cohesin protein-depletion and found that only at several hundred loci, newly gained enhancers-promoters (E-P) loops can be reproducibly observed upon CTCF depletion. The new E-P loops are cohesin-dependent and associated with recurrent gene activation. These findings allow us to map FINs and define their direct target genes genome-wide. FINs are mostly present in euchromatin, do not overlap with TAD boundaries, and can be perturbed by *Wapl*-depletion or CTCF-displacement. We therefore conclude that FINs, but not TAD boundaries, are *bona fide* insulators.

## Introduction

It is generally accepted that TADs confine the communication between genes and enhancers for transcriptional control^1, 2, 3, 4, 5, 6^. Most of the TADs are maintained by CTCF and cohesin, which are also known as insulator proteins. Cohesin is a multimeric complex consisting of *SMC1A*, *SMC3*, *RAD21* and one SA subunit (*STAG1* or *STAG2*). TAD forms when a DNA loop extrudes out of the cohesin complex ring before blocked by CTCF binding sites in convergent orientation, and cohesin serves as an enzymatic engine for this process. Deletion of CTCF or cohesin core subunit RAD21 leads to near complete loss of TADs^7, 8^. Cohesin function is also regulated by cohesin loading complex MAU2/NIPBL^9^ and cohesin release factor WAPL^10, 11, 12^: TADs disappear upon loss of NIPBL^13, 14^, while loss of WAPL causes bigger loops because cohesin can pass through CTCF blocks due to prolonged loop extrusion^14, 15^.

Insulators are canonically defined as the DNA elements that inhibit gene expression when put between enhancers and promoters^16, 17, 18^. The discovery of thousands of CTCF-demarcated TADs immediately suggests a model that CTCF sites at TAD boundaries are enhancer-blocking insulators, which is supported by genetic experiments focusing on selected loci^19, 20, 21, 22, 23^.

However, this model is challenged by a genome-wide Hi-C study in mESCs (Nora *et al.*, **Fig. 1a**) which only observed mild transcription changes despite near complete loss of TADs upon CTCF-depletion^7^. This conclusion is reproduced in two other genome-wide studies done with *in situ* Hi-C and micro-C^24, 25^ (**Fig. 1a**). Notably, Kubo *et al*. reported a role of CTCF in mediating E-P interactions^24^, and Hsieh *et al.* reported short-range cohesin-independent cross-TAD E-P interactions^25^. Taken together, although all these studies questioned the insulating function of TAD boundaries, they did not address the fundamental question about the whereabouts of the real enhancer-blocking insulators.

**Figure 1.**
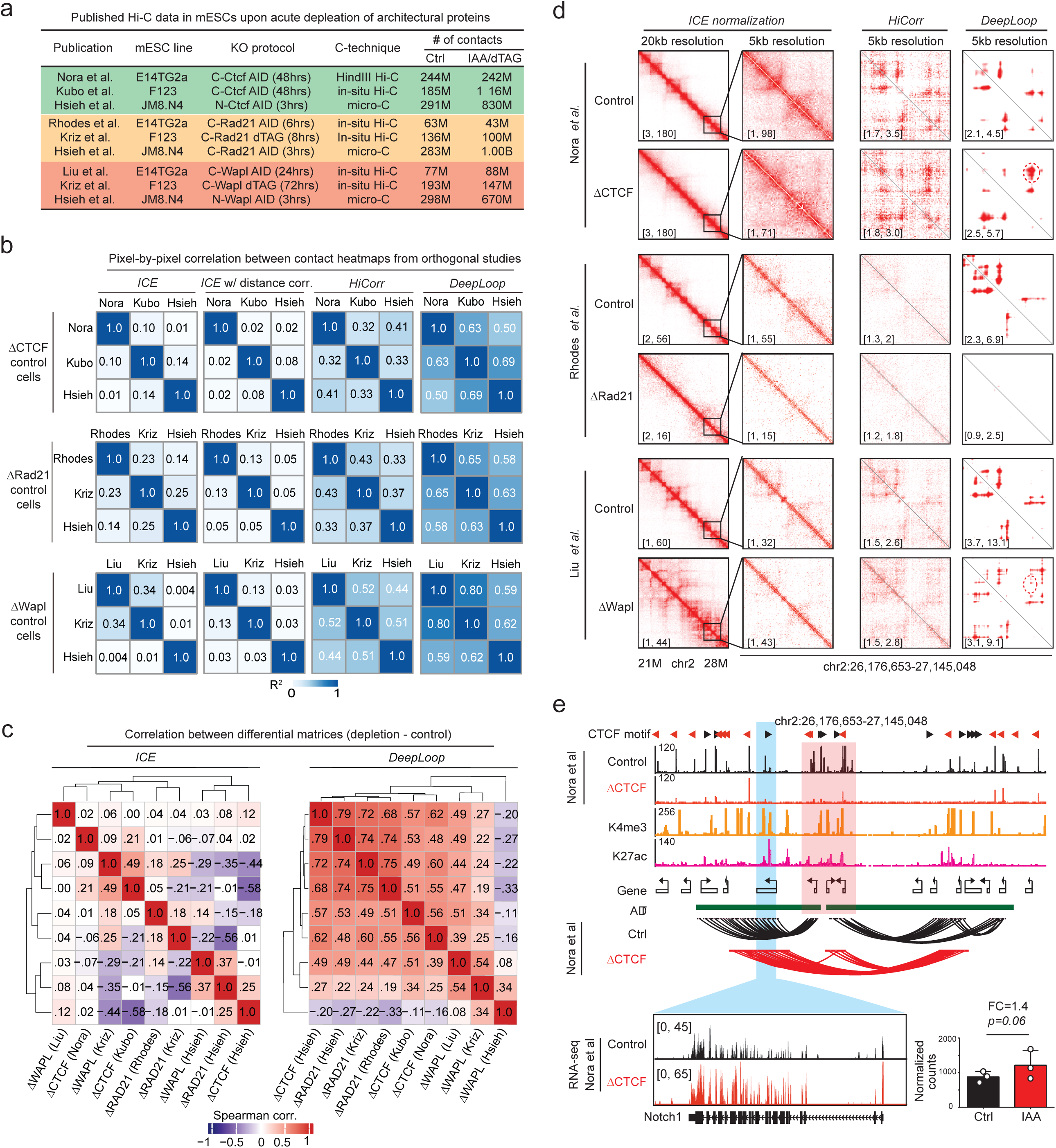
*DeepLoop* allows cross-platform comparison between independent CTCF- and cohesin-degron Hi-C datasets at loop-resolution. **a,** A list of published Hi-C and micro-C studies with acute depletion of *Ctcf*, *Rad21*, or *Wapl* in mESCs. **b,** Matrices summarizing R-square values when comparing contact heatmaps from different pipelines and orthogonal Hi-C/micro-C datasets in control mESCs. **c,** Correlation matrices comparing the differential matrices of all nine degron experiments. For each degron experiment, the differential matrix is the difference between two Hi-C contact matrices (depletion – control) at 5kb resolution. Left, Hi-C contact matrices from *ICE* are used to compute differential matrices; right, Hi-C contact matrices from *DeepLoop* are used. **d,** Contact heatmaps of the CTCF-, Rad21-, and Wapl-degron Hi-C data at *Notch1* locus with *ICE*, *HiCorr* and *DeepLoop*. Red dashed circle: gained loops. Numbers in brackets indicate the color scale of heatmaps: for *ICE* normalization and *HiCorr* heatmaps, the maximum intensity corresponds to the 98^th^ percentile of the corresponding contact matrix; for *DeepLoop* heatmaps, the maximum intensity corresponds to the lower cutoff of top 500,000 pixels genome-wide. **e**, Genome browser tracks showing ChIP-seq data and Hi-C loops from Nora *et al.* at *Notch1* locus. Blue highlighted region showing Notch1 gene which is at the anchor of a gained loop. Pink region highlights the potential insulator region that blocks the formation of *Notch1* E-P loop. RNA-seq results are also shown for *Notch1* gene in WT (black) and CTCF-depleted cells (red). Bar plot: quantitation of *Notch1* expression from RNA-seq. P-value: one-side Wald test provided by DE-seq2.

In this study we demonstrated that “functional insulators” (FINs) can be identified by examining if their flanking regions gain E-P interactions upon CTCF depletion. Challenges exist to map FINs with Hi-C-like genome-wide technologies: (i) most of the public Hi-C datasets do not have billion-scale read depth usually required for loop-level analysis (**Fig. 1a**); (ii) E-P interactions are generally weaker than CTCF loops in Hi-C and therefore harder to detect; (iii) the gain of E-P loop signal can be subtle, requiring robust quantitation of loop strength; (iv) it is difficult to rule out the possibility that some loop changes are secondary effects after prolonged CTCF or cohesin depletion^7, 25^. These challenges can be addressed by comparing independent datasets, but existing CTCF-depletion data are from different Hi-C protocols, which perform very differently in loop-level analyses with conventional pipelines^26^. We previously developed *HiCorr* and *DeepLoop* pipelines to allow robust comparison of loop profiles from orthogonal Hi-C and micro-C platforms, even when sequencing depth is low^27, 28^. Here we use *DeepLoop* to perform a meta-analysis of orthogonal Hi-C datasets to map recurrent DNA rewiring events upon depletion of CTCF and other cohesin proteins (**Fig. 1a**), thus generating a reliable genome-wide map of enhancer-blocking insulators.

## Results

### *DeepLoop* allows cross-platform comparison of independent CTCF- and cohesin-degron Hi-C datasets at loop-resolution

We collected nine pairs of independent Hi-C or micro-C datasets from six publications in which CTCF^7, 24, 25^, RAD21^25, 29, 30^, or WAPL^25, 29, 31^ were acutely depleted in three different mESC lines. The number of contacts in these Hi-C/micro-C datasets are highly variable ranging from 43 million to ∼1 billion. For every target protein, the choice of degron methods (AID or dTAG) and the timespan of protein degradation (range from 3hr to 3 days) are also variable (**Fig. 1a**).

To assess how these variations complicate cross-platform comparisons. We first performed low resolution compartment- and TAD-level analysis on the three CTCF-degron datasets with conventional pipelines. At compartment level (500kb resolution), all datasets display rather consistent compartment correlation heatmaps, and the compartmental organization is largely preserved after CTCF depletion (**Supplementary Fig. 1a-c**). When plotting ICE-normalized contact matrices^32^ at 40kb resolution followed by DI (directionality index)-based TAD analyses, all three studies consistently show similar TAD structures and a massive loss of TADs upon CTCF depletion despite a trend that micro-C detected more bins with stronger directionality biases (**Supplementary Fig. 1d-f**). These observations confirmed the findings from the original reports and showed that conventional pipelines are overall robust across different Hi-C platforms at low-resolution.

However, Hi-C analysis at high resolution is severely affected by data sparsity and protocol biases^33, 34^, thus the robustness of cross-platform comparison is highly dependent on the choice of analytical pipelines. We generated 5kb resolution contact heatmaps (each pixel in a heatmap represents a pair of 5kb bins) using conventional *ICE*^32^ (with or without distance correction), *HiCorr*^34^, and *DeepLoop*^28^, then used scatter plots to compare control cell heatmaps from independent experiments to assess the consistency of different pipelines in analyzing orthogonal Hi-C platforms (**Supplementary Fig. 2**). Here we only compare control cell heatmaps because before protein depletion, the control cells from all studies are expected to be at similar biological states. Protein-depletion cells from independent studies, even if they depleted the same protein, can be still biologically diverse due to many variations such as time-span of protein depletion, incomplete protein depletion, loss of stemness, *etc.* We found that although conventional ICE pipelines show poor cross-platform consistency with R-square values often close to zero, *HiCorr* and *DeepLoop* clearly improved the consistency with much higher cross-platform R-square values (**Supplementary Fig. 2**, summarized in **Fig. 1b**).

We also use scatter plots to compare the heatmaps before and after protein depletion at pixel-by-pixel level and interestingly, found that only HiCorr and DeepLoop heatmaps show recognizable populations of loop pixels that retain their strength upon CTCF- or Rad21-depletion (**Supplementary Fig. 2**), suggesting their robustness in revealing the unchanged loop pixels that are masked by noises in the other pipelines. To further evaluate the cross-platform consistency when applying different pipelines to analyze dynamic loops, we generated differential heatmaps by subtracting each pair of 5kb-resolution contact heatmaps (depletion – control), then computed the correlations between the differential heatmaps from all nine pairs of degron experiments (**Fig. 1c**, **Supplementary Fig. 3**). Only the matrices from *HiCorr* and *DeepLoop* clustered CTCF- and Rad21-degron datasets together and *Wapl*-degron datasets are separated out (**Fig. 1c**, **Supplementary Fig. 3**). This is consistent with the knowledge that depletion of *CTCF* or Rad21, but not Wapl, causes massive loss of TADs and CTCF loops. Taking together, these results support the robustness of *DeepLoop* in cross-platform analysis regardless of read depth^28^.

### Hi-C meta-analysis reveals recurrent E-P loop-gaining events upon CTCF depletion

We first focused on the CTCF-degron Hi-C datasets. In the example of **Fig. 1d** (top two rows), with conventional ICE-normalization, we observed dampened TAD structure at 20kb resolution as the original study reported^7^, but at 5kb resolution, loop signal from ICE-normalized heatmaps are weak and often invisible. From the *DeepLoop* enhanced heatmaps, we can consistently observe multiple CTCF loops that appear in all control cells heatmaps but are substantially weakened upon CTCF depletion. A notable change in this region is the gain of long-range loops (**Fig. 1d**, second row, loop pixels in dashed circle) connecting *Notch1* gene to a distal enhancer region with multiple H3K27ac peaks (**Fig. 1e**). Consistent with gain of E-P loop, *Notch1* is upregulated upon CTCF-depletion (**Fig. 1e**). The gain of loop signal at this locus is also observable in *Wapl*-depleted cells (**Fig. 1d**, bottom two rows) and two other CTCF-degron studies (**Supp. Fig. 4**), but not in the *Rad21*-depleted cells (**Fig. 1d**, 3^rd^ and 4^th^ rows, **Supp. Fig. 4g**). These observations suggest that the two CTCF loop anchors at the TAD boundary region (**Fig. 1d**, highlighted in pink) are blocking the *Notch1*-enhancer interaction. This example also demonstrated that *DeepLoop* may detect recurrent insulating events from CTCF-degron Hi-C data without the need for ultra-high sequencing depth.

To perform a genome-wide analysis of insulating events, we call 300K∼500K loop pixels from each CTCF-degron dataset and use the pixel intensity in *DeepLoop* heatmaps to measure the loop strength (**Fig. 2a, Supp. Fig. 5**). Notably, *DeepLoop* outputs are enhanced from ratio (obs/exp) heatmaps instead of normalized contact frequency used by ICE/KR. We previously showed that this approach promises cross-platform convergence between orthogonal Hi-C and micro-C data^28^.

**Figure 2.**
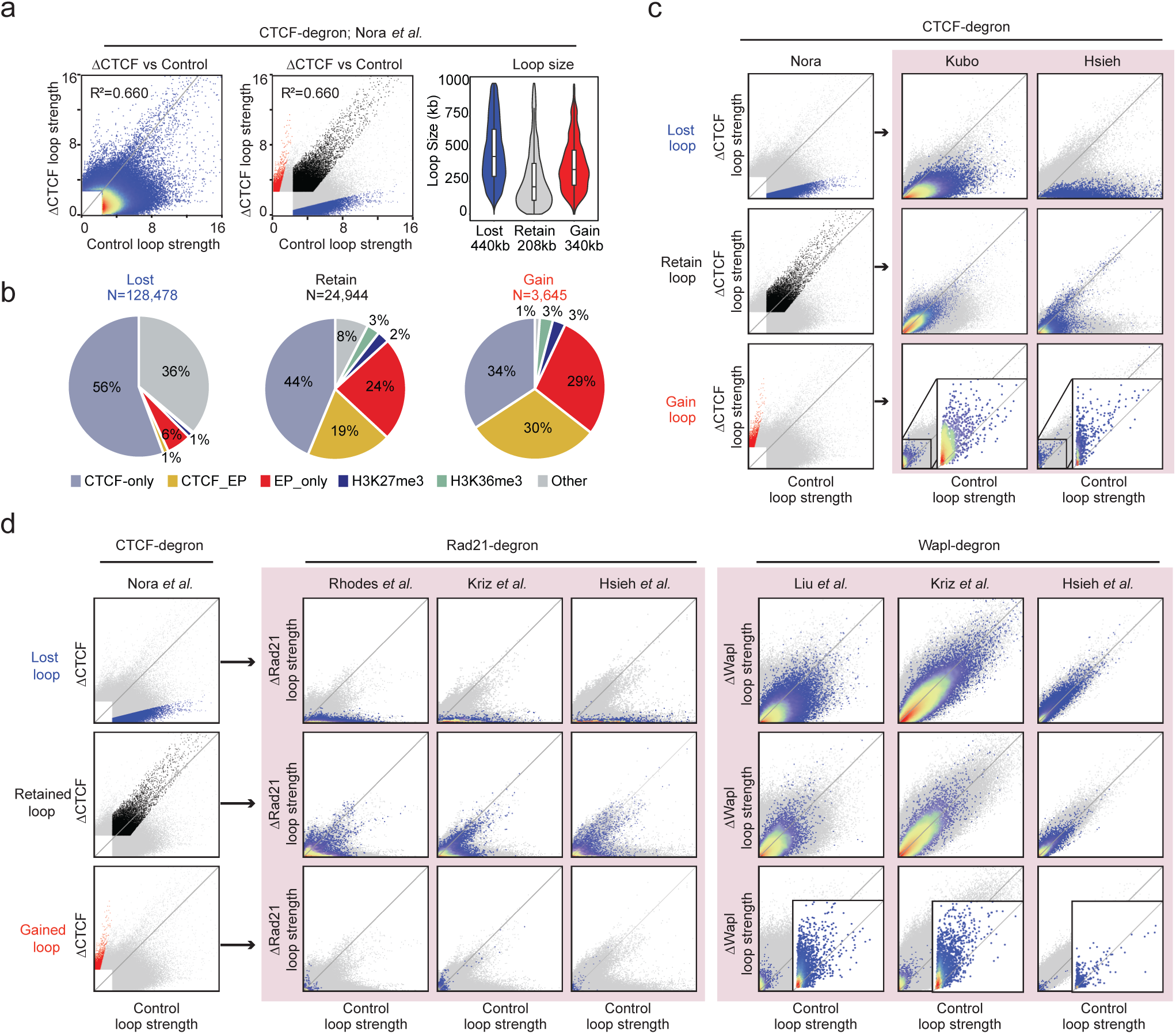
Identify and characterize recurrent gain of E-P loop events in CTCF-depleted cells through cross-platform Hi-C meta-analyses. **a-c,** Meta-analysis of independent CTCF-degron Hi-C datasets. **a**, Left: scatterplot comparing the strength of loop pixels between WT and CTCF-depleted mESCs (every dots represent one pixel in the contact heatmap); middle: the same scatterplot as left but highlight the lost (blue), retained (black), and gained (red) loop pixels; right: violin plot comparing the size of loop pixels and the value below each plot indicates the median loop size for the corresponding category. **b,** Pie charts showing the components of lost, retained and gained loop pixels based on epigenetic marks. **c**, Scatter plots showing the consistency of lost, retained and gained loop pixels between three CTCF degron studies. Plots in the left column highlight the lost, retained, and gained loop pixels from Nora *et al*., the right two columns show the quantitative changes of the corresponding pixels in *Kubo et al.* or *Hsieh et al*. **d,** The lost, retained, and gained loop pixels from one CTCF-degron Hi-C data (Nora *et al.*) are plotted onto Hi-C scatter plots from *Rad21*-or *Wapl*-degron studies to examine their quantitative changes.

In all CTCF-degron studies, the scatter plots are clearly down-skewed reflecting a massive loss of chromatin loops upon CTCF deletion (**Fig. 2a, Supp. Fig. 5a, d**, left panels). Using a simple fold-change cutoff, we called loop pixels that lose strength (“lost”), retain strength (“retained”), or gain strength (“gained”) from each dataset (**Fig. 2a, Supp. Fig. 5a, d**, middle panels). We further classified the loop pixels based on their epigenetic marks (**Methods**, examples of loop classification in **Supp. Fig. 6**). “CTCF-only” loop pixels, defined as those with at least one anchor overlapping a CTCF peak, were predominantly found among the “lost” pixels. In comparison, “retained” and “gained” loop pixels are enriched with “CTCF-EP” or “EP-only” loops characterized by promoter (H3K4me3) or enhancer (H3K27ac) marks at one or both anchors (**Fig. 2b, Supp Fig. 5b, e**). Interestingly, in all three studies the “gained” loops are much larger than the “retained” loops (**Fig. 2a, Supp. Fig. 5a, d**, right panels); we hypothesized that this is because the “gained” E-P loops form through cohesin loop extrusion (more discussions below).

We also examine the robustness of the “lost”, “retained”, and “gained” pixels across independent Hi-C/micro-C studies. As expected, the “lost” pixels from any study generally also lose strength in the other two studies (**Fig. 2c, Supp Fig. 5c,f**, first row). However, the consistency of “retained” loop pixels is study specific: (i) although the “retained” loop pixels from Nora *et al.* are mostly also “retained” in Kubo *et al.*, some of them lose strength upon CTCF degradation in Hsieh *et al.* (**Fig. 2c**, second row); (ii) most of the “retained” loop pixels from Kubo *et al*. lose strength in the other two studies (**Supp. Fig. 5c**, second row); (iii) the “retained” loops from Hsieh *et al.* remain “retained” in both Nora *et al.* and Kubo *et al.* (**Supp. Fig. 5f**, second row). Since incomplete CTCF depletion causes the leftover CTCF loops^7^, these observations suggest that CTCF depletion in Kubo *et al.* may be less complete than the other two studies. On the other hand, Hsieh *et al.* appear to have achieved better CTCF depletion, perhaps due to its use of an N-terminal degron system (**Fig. 1a**), although platform-specific biases, differences in assay sensitivity and sequencing depth may also explain some of the discrepancies between studies.

Interestingly, the “gained” pixels from any one of the three studies generally also gain strength in the other two studies (**Fig. 2c, Supp Fig. 5c,f**, third row). The only exception is that some “gained” loop pixels from Kubo *et al*. lose strength in Hsieh *et al*. (**Supp Fig. 5c**). Again, this is likely because Kubo *et al.* retains more CTCF loops due to incomplete CTCF degradation.

Nevertheless, most of the “gained” pixels from Nora *et al*. and Hsieh *et al*. do gain strength in Kubo *et al.* (**Fig. 2c, Supp Fig. 5f**), supporting the consistency between the three studies.

Notably, E-P loop gaining events are largely neglected in the original Hi-C/micro-C studies because the changes are often too subtle to distinguish from background with conventional analytical tool (*e.g.*, **Fig. 1d**). Our meta-analyses prove that although there are only a small number of newly gained E-P loops, they are reliable consequences of CTCF-depletion; affected genes such as *Notch1* in **Fig. 1** are putative target genes of CTCF-mediated insulation.

### Hi-C Meta-analyses of *Rad21*- and *Wapl*-depletion data distinguish consensus loop regulatory mechanisms from system variations

We also performed meta-analyses of the *Rad21*- and *Wapl*-depletion Hi-C datasets. Ablation of *Rad21* also causes massive loss of CTCF loops, while the “retained” and “gained” loops are also enriched with E-P loops (**Supp. Fig. 7a-i**). However, there are important differences between *Rad21*- and *CTCF*-depleted cells. (i) In *Rad21*-depleted cells, the “retained” and “gained” loop pixels (except the “gained” loop pixels in Rhodes *et al.*) are much smaller (mostly within 100kb) than those in *CTCF*-depleted cells, indicating that cohesin-independent loops are short-range (**Supp. Fig. 7a, d, g**, right panels). One alternative explanation is that the greatly reduced pool of cohesin only allows rare and short-lived extrusion events, resulting in smaller loops. (ii) Unlike in CTCF-depleted cells, the “gained” loop pixels are NOT recurrent between the three *Rad21*-degon datasets (**Supp. Fig. 7c, f, i**, last row). (iii) The “retained” and “gained” loop pixels in *Rad21*-depleted cells (except the “gained” loop pixels in Rhodes *et al.*) are more enriched with H3K36me3 mark, suggesting transcription associated chromatin interactions (**Supp. Fig. 7b, e, h**).

The “gained” loops in *Rad21*-depleted cells from Rhodes *et al*. are enriched with H3K27me3 and tends to be larger in size (**Supp. Fig. 7a, b**, right panels). These observations are consistent with the conclusion from the original study that cohesin removal enhances long-range interactions between a subset of polycomb-occupied regions^30^. However, these features (the larger loop size and the enrichment with H3K27me3) are specific to “gained” loops in Rhodes *et al.* but not observed in the other two *Rad21*-degron studies (**Supp. Fig. 7d, e, g, h**, right panels). We manually examined the “gained” H3K27me3 loops in Rhodes *et al.* and found that in almost all cases, the long-range H3K27me3 loops are already present in the control cells from Kriz *et al.* and Hsieh *et al.* before *Rad21* depletion (two examples are shown in **Supp. Fig. 8**). Therefore, at least in the control mESCs from Kriz *et al.* and Hsieh *et al.*, cohesin does not prevent the formation of long-range H3K27me3 loops.

In contrast, ablation of cohesin unloading factor *Wapl* generates up-skewed scatterplots in all three studies (**Supp. Fig. 7j, m, p**, left panels). The skewness of Hsieh *et al.* is less apparent, probably because Hsieh *et al.* performed *Wapl* depletion for much shorter time (3hrs) than Liu *et al*. (24hrs) and Kriz *et al*. (72hrs) (**Fig. 1a**). In all *Wapl*-degron studies, the “lost”, “retained”, and “gained” loops are almost equally enriched with CTCF and E-P loops (**Supp. Fig. 7k, n, q**).

Consistent with the model that *Wapl*-depletion stabilizes chromatin-bound cohesin and promote loop extrusion^14, 15^, the “gained” loops are significantly bigger than “lost” and “retained” loops in all three studies (**Supp. Fig. 7j, m, p**, right panels). The “lost”, “retained”, and “gained” loop pixels are generally consistent between studies (**Supp. Fig. 7l, o, r**) despite some discrepancy for “lost” loop pixels from Hsieh *et al.* (**Supp. Fig. 7r**), again likely because Hsieh *et al.* only depleted *Wapl* for a short period of time.

### Newly gained E-P loops in CTCF-depleted cells are cohesin dependent

We next take the “lost”, “retained”, and “gained” loop pixels from *CTCF*-degron studies and examine how they change in *Rad21*- and *Wapl*-degron data. Firstly, nearly all “lost” loops in *CTCF*-depleted cells also disappear in *Rad21*-depleted cells (**Fig. 2d**, first row, **Supp Fig. 9a,b**, first row), suggesting that most of the CTCF loops are cohesin-dependence. Secondly, a subset of the “retained” loops in CTCF-depleted cells are also “retained” upon *Rad21*-depletion (**Fig. 2d**, second row, **Supp Fig. 9a, b**, second row), suggesting cohesin-independence. The other “retained” loops in CTCF-depleted cells lose strength upon *Rad21*-depletion likely because they are: (i) leftover CTCF loops due to incomplete CTCF-depletion, or (ii) non-CTCF loops formed via cohesin-dependent mechanisms. Interestingly, most of the “retained” loops in CTCF-depleted cells from Kubo *et al.* are lost upon *Rad21*-depletion (**Supp Fig. 9a**, second row), but most of the “retained” loops from Hsieh *et al.* are also retained upon *Rad21*-depletion (**Supp Fig. 9b**, second row), again suggesting that Hsieh *et al.* may have achieved more complete CTCF-depletion than Kubo *et al*.

Finally, the “gained” loops in CTCF-degron studies, which are weak in both CTCF-degron and Rad21-degron control mESCs, do not gain strength upon *Rad21*-depletion cells (**Fig. 2d, Supp Fig. 9a, b**, third row), suggesting that the formation of these “gained” loops requires cohesin. In contrast, we observed that these “gained” loops in CTCF-depleted cells showed an apparent upward bias in the scatter plots of *Wapl*-degron studies (**Fig. 2d, Supp Fig. 9a, b**, third row).

This observation is consistent with the example in **Fig. 1d** and further supports a role of cohesin in forming those “gained” loops. *Wapl*-depletion may perturb the function of insulators by releasing short-range CTCF loops and allow insulated E-P interactions to skip some insulators.

### Define, map, and characterize functional insulators (FINs) in mESC genome

Among the total 8,141 “gained” loop pixels from all three CTCF-degron Hi-C/micro-C datasets, 4,944 pixels are recurrent in at least two studies with a median size ∼250kb (**Fig. 3a**). In each dataset, the “gained” loop pixels can be merged into ∼300 “loop-gaining loci” (**Methods**) and there are substantial overlaps between independent studies: 174 loop-gaining loci are shared by all three studies and 304 are shared by at least two studies (**Fig. 3b**). We hypothesize that these recurrent E-P loop-gaining loci are caused by loss of CTCF insulation; indeed, 4,665 out of the 4,944 loop-gaining pixels (94%) are insulated by at least one CTCF loop in wildtype cells (**Fig. 3c**). We further curated the reproducible loop-gaining loci based on the types and numbers of CREs involved (**Fig. 3d**). Nearly all the curated loci (283 of 299) gain E-P loops (2 loci gain CTCF contacts and the others cannot be clearly classified). We further classified the E-P loop-gaining loci into three categories (**Fig. 3d**). In *Type I*, one of the two E/P sites is insulated in a CTCF loop while the other site is not (exampled in **Supp. Fig. 10a**); *Type II* involves at least two CTCF loops insulating two E/P-sites (exampled in **Fig. 1e, Supp. Fig. 10b**). In contrast, the gained E-P loops in *Type III* do not have insulating CTCF loops (exampled in **Supp. Fig. 10c**). *Type I* and *II* represents most of the E-P loop gaining loci (263 out of 283), while only 20 loci belong to *Type III*, again suggesting that most of the loop gaining loci are due to the removal of CTCF insulation.

**Figure 3.**
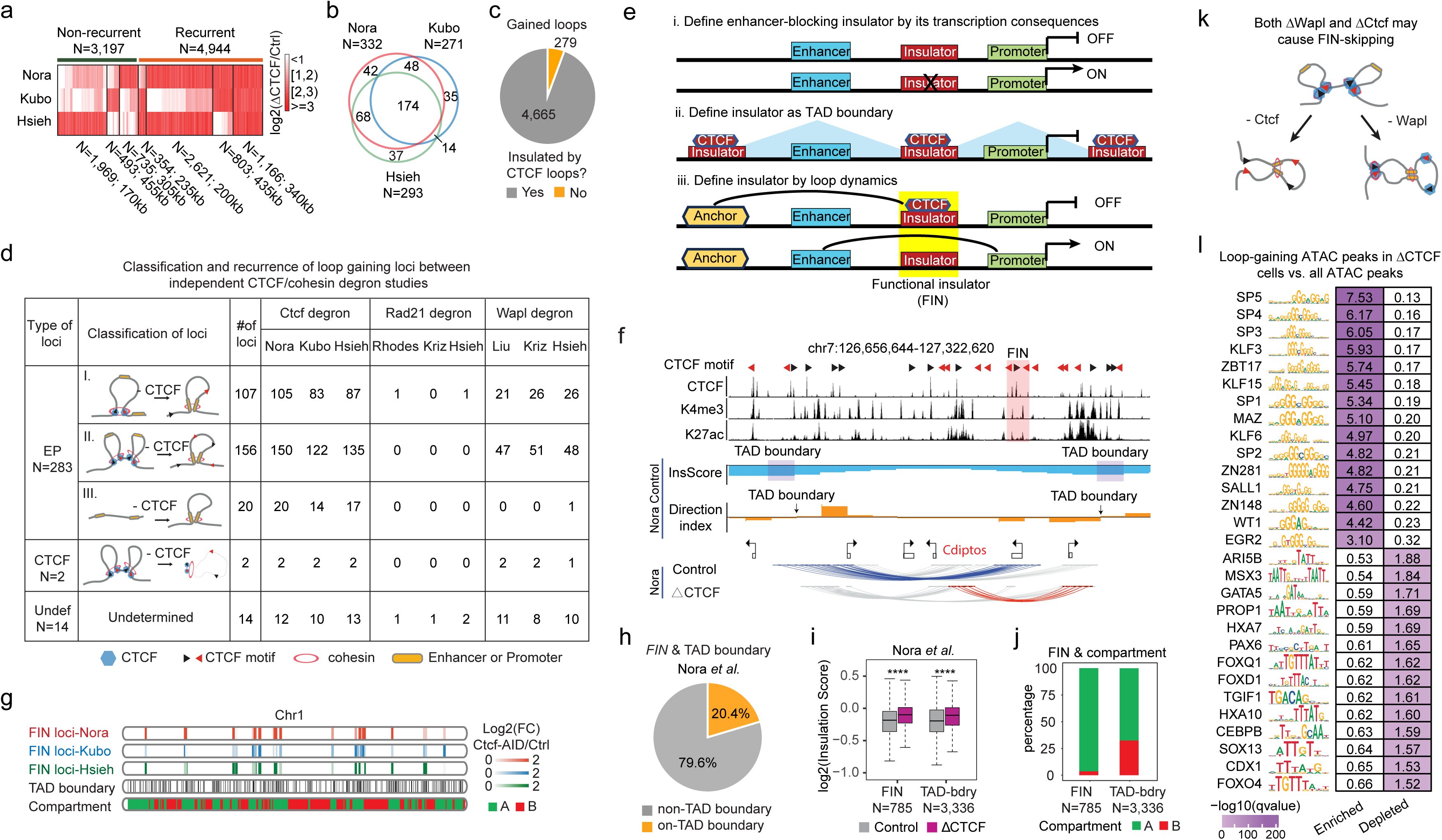
Characterization of functional insulators (FINs) in mESC genome. **a,** Heatmap showing the union of all gained loop pixels from three CTCF-degron Hi-C datasets. Every row represents one CTCF-degron study, and every column represents one loop pixel. Loop pixels are organized into clusters based on their reproducibility. The number of loop pixels in each cluster and their median size are also shown. Color scale indicates the increase of loop signal in log scale. **b**, Venn diagram showing the consistence of loop-gaining loci identified from three datasets. **c,** Pie chart showing that most of the gained loops in CTCF-depleted cells have at least one CTCF loop anchor in between in wildtype cells. **d,** A table summarizing the classification and recurrence of loop gaining loci between independent CTCF- or _ohesion-degron studies. **e,** Cartoons summarizing three possible ways to define insulators. **f,** An example of non-TAD boundary FIN. The FIN is highlighted in pink. The tracks of InsScore and DI are also included. TAD boundaries determined from InsScore and DI are highlighted in purple and indicated with black arrows, respectively. **g,** Ideogram showing the locations of loop-gaining loci from three studies, TAD boundaries, and compartments on chromosome 1. FINs from three different CTCF-degron studies are shown in different colors, and the color scale shows the fold increase of gained loop in log scales. **h,** Pie chart showing the percentage of FINs that overlap TAD boundary. **i,** Boxplot comparing the insulation scores at FINs and TAD boundaries before and after CTCF depletion computed with HI-C data from Nora *et al*. **h,** Stack bar plot comparing the proportion of FINs and TAD boundaries in compartment A or B. **I,** A model to explain why some of the newly gained E-P loops in CTCF-depleted cells are also observed in Wapl-depleted cells. **j,** A list of TF motifs enriched or depleted in the mESC ATAC-seq peaks that will gain chromatin loops upon CTCF depletion. Enrichment or depletion are relative to all mESC ATAC-seq peaks.

We therefore propose to define “functional insulators” (FINs) as the CTCF binding sites within the “lost” loop anchors that when bound by CTCF, can actively block the interactions between flanking *cis*-regulatory elements (CREs) (model *iii* in **Fig. 3e, Methods**). Therefore, FINs can be found from “loop-gaining loci” and in fact, most “loop-gaining loci” contain at least one FIN, and we therefore also call them “FIN loci” (**Methods**). The definition of FIN is based on the gain-of-loop upon loss of insulator function, which is conceptually analogous to the conventional model to define insulators based on the gain-of-expression upon loss of insulator function (model *i* in **Fig. 3e**) but distinct from the TAD-based definition of insulators (model *ii* in **Fig. 3e**), which is a static definition that does not need the knowledge about the outcomes after loss of insulators.

In **Fig. 3f**, we show a FIN (highlighted in pink) that is not at TAD boundary. In this example we also included the browser tracks for “insulation score” (InsScore)^35^ and “directionality index” (DI)^1^. TAD boundaries can be defined as bins where InsScore reaches local minimum or positions where DI changes its sign (**Supp. Fig. 11a**), but the FIN in this example does not fit into either criterion. There are much fewer FINs than TAD boundaries (**Fig. 3g**) and only a small percentage of the FINs overlap with TAD boundaries (**Fig. 3h, Supp. Fig. 11b**). These observations indicate that most FINs are inside TADs and conversely, loss of TAD boundaries usually does not cause loop formation across their flanking regions. Change of InsScore also does not predict FINs because although we observed an overall trend of increased InsScore at FINs upon CTCF depletion, TAD boundaries also show the same trend and cannot be distinguished from FINs (**Fig. 3i, Supp. Fig. 11c**). Interestingly, most (96%) of the FINs are within compartment A (**Fig. 3j**), indicating that active E-P blocking mainly happens within euchromatin region. Regardless, it is important to note that low-resolution 3D genome features, including compartment and TAD boundaries, lack the precision to reveal the dynamic changes of loop profiles as we used to define FINs.

Cohesin degron Hi-C data further supports a cohesin-dependent mechanism of FINs. We found that almost none of the curated loop-gaining events appear in the *Rad21*-degron cells (**Fig. 3d**), indicating cohesin dependency. Interestingly, a significant fraction of the *Type I* and *II*, but not the *Type III* “gain of E-P loop” loci, can be also observed in *Wapl*-degron datasets (**Fig. 3d**).

This phenomenon can be explained by previous findings that *Wapl* deletion allows loop extrusion to skip CTCF blocks^14, 15^. As shown in the carton in **Fig. 3k**, because *Wapl*-depletion stabilizes chromatin-bound cohesin and leads to longer loop extrusion, previously insulated enhancers and promoters can interact inside the bigger CTCF loop due to “insulator skipping”, creating the same E-P loop as observed in CTCF-depleted cells. All these results support FINs as *bona fide* insulators of cohesin-mediated loop extrusion.

To explore which types of enhancers or promoters would gain chromatin loops upon CTCF depletion, we picked the FIN-blocked ATAC-peaks in mESCs (n = 995) and scan for enriched or depleted motifs compared to all mESC ATAC-peaks (n = 59,871). Interestingly, the loop-gaining ATAC-peaks are enriched with many G-rich motifs but depleted with A/T-rich motifs (**Fig. 3l**).

This result is reminiscent of previous reports that G-quadruplexes (G4) structure may regulate chromatin looping^36, 37^. Notably, many of the putative transcription factors binding to these G-rich motifs have been reported to regulate loop formation in various cell types, such as Kruppel-like factors^38, 39, 40^, SP1^41^, MAZ and several zinc-finger proteins^42, 43, 44^. Therefore, our results suggest that G-rich *cis*-regulatory elements are more likely to act as new loop anchors upon CTCF-depletion.

### FINs are associated with consistent early transcription responses upon CTCF depletion

We found that in all CTCF-degron studies FIN proximal regions (+/- 100kb) are 2∼3 fold more enriched with DEGs than TAD-boundary proximal regions, and the “gained” loop anchors are 10 times more enriched with DEGs (**Fig. 4a**). Furthermore, early response DEGs are clearly more enriched at gained loop anchors (**Fig. 4b**), suggesting that FIN-blocked enhancers and promoters are the direct targets of insulators. In 16.8% of FIN loci (49 out of 299), we observed the blocking of 43 super-enhancers (there are 231 super-enhancers in total) (**Fig. 4c**). As expected, genes gaining super-enhancer interactions are more likely to be upregulated than genes gaining interactions with regular enhancers (**Fig. 4d**).

**Figure 4.**
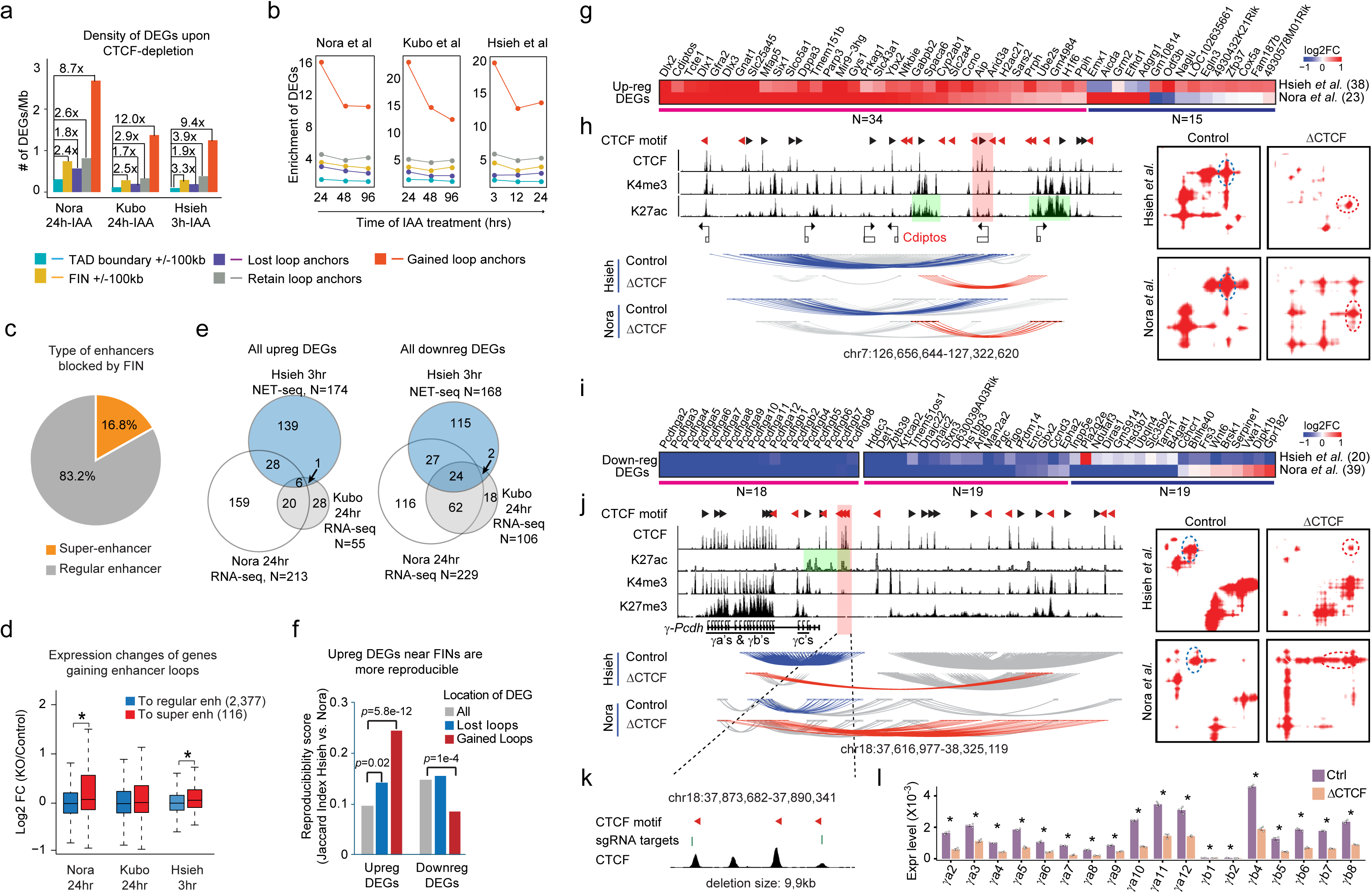
FINs are associated with consistent early transcriptional consequences in independent CTCF-degron studies. **a,** Bar plots comparing the density of DEGs upon CTCF-depletion across different genomic regions. The y-axis indicates the number of DEGs per Mb. The number of DEGs were calculated from the following datasets: 3-hour NET-seq for Hsieh *et al.*, 24-hour RNA-seq for Nora *et al*., and 24-hour RNA-seq for Kubo *et al.* **b,** Enrichment of DEGs at different types of genomic regions compared to whole genome as background. X-axis: time of IAA treatment (hrs) in each study; y-axis: fold enrichment. Each panel represents one CTCF-degron study. **c,** Pie chart showing the percentage of FINs that are blocking super enhancers; **d,** Boxplot showing the expression changes of genes gaining enhancer loop. Red: genes gaining super-enhancer loop; Blue: genes gaining regular enhancer loop. *, p < 0.05, Wilcoxon signed-rank test. **e,** Venn diagrams showing the overlap of all up- (left panel) and down-regulated (right panel) DEGs between three CTCF-degron studies. **f,** Bar plot comparing the reproducibility scores (measured by Jaccard Index) of up- or down-regulated DEGs between two CTCF-degron studies (Hsieh *et al.* and Nora *et al.*). Gray bars: all DEGs; blue bars: DEGs at the anchors of lost loops; red bars: DEGs at the anchors of gained loops. P-values are from binomial test. **g,** Heatmap showing the union of upregulated DEGs at the anchors of gained loops from *Hsieh et al* and *Nora et al*.; DEGs are clustered based on consistency. **h,** Left panel shows the browser tracks near *Cdiptos* locus and right panel shows the Hi-C heatmaps in the same region from *Hsieh et al* and *Nora et al.* The direction of CTCF motifs are indicated by triangles. Blue arches (left) and blue dashed circles (right): lost loop pixels; red arches and red dashed circles: gained loop. Regions highlighted in green: enhancers that gained loops; the region highlighted in pink: a CTCF loop anchor is defined as *FIN*. **i,** Similar as **g**, a heatmap showing the union of down-regulated DEGs at the anchors of gained loops from *Hsieh et al.* and *Nora et al*. Among these genes 18 genes are from the Pcdh-γ locus (shown as a separate heatmap). **j,** Left panel shows the browser tracks near Pcdh-γ locus and right panel shows the Hi-C heatmaps from two CTCF-degron studies. **k,** Design of sgRNAs for insulator deletion. **l,** RT-qPCR shows the down-regulation of Pcdh-γ genes in mESCs after insulator deletion.

Using common cutoffs (2-fold for RNA-seq and 1.5-fold for NET-seq, p<0.05), each of the CTCF-degron studies identified a few hundred early response DEGs, and the reproducibility between independent studies is limited due to complex reasons (**Fig. 4e**). Both Nora *et al.* and Kubo *et al.* performed RNA-seq and the earliest timepoint was 24hr CTCF-depletion, which may still include late-response DEGs caused by indirect mechanisms (*e.g.*, loss of pluripotency). Among the two RNA-seq studies, Kubo *et al*. detected fewer DEGs than Nora *et al.*, but 70% of Kubo *et al*.’s DEGs are also observed in Nora *et al.* (47% of the upregulated DEGs and 81% of the downregulated DEGs, **Fig. 4e**). This observation is consistent with our discussion above that CTCF depletion in Kubo *et al*. was less complete. Hsieh *et al*. performed both NET-seq and RNA-seq after only 3hr CTCF-depletion, which should include fewer late-response genes, but the short treatment time may also reduce the sensitivity of DEG detection. We use NET-seq data because Hsieh *et al.* reported that NET-seq has higher sensitivity in identifying DEGs^25^ as it maps newly synthesized RNA, while RNA-seq measures the amount of steady state RNA in the cells.

We decided to focus on the comparison between Hsieh *et al.* and Nora *et al.* as they represent a more complete set of CTCF direct targets. We next hypothesize that DEGs near FINs are more likely to be the direct targets of CTCF-mediated enhancer-promoter insulation. If this hypothesis is true, DEGs near FINs should be more reproducible between independent studies. Indeed, the upregulated DEGs near FINs (*i.e.*, at “gained” loop anchors upon CTCF depletion) are more reproducible, but downregulated DEGs near FINs are less reproducible than controls (**Fig. 4f**, red bars). In contrast, the “lost” loop anchors does not predict the reproducibility of DEGs. This observation suggests that transcription activation is the primary consequence of insulator loss, consistent with the convention concept that insulators block enhancer-mediated gene activation.

In **Fig. 4g** we plotted 49 upregulated DEGs at FIN loci as putative “direct targets” of insulators (from Hsieh *et al.* or Nora *et al.*), among them 34 are considered as “high-confidence” targets supported by both studies. One representative example is shown in **Fig. 4h** and **Supp. Fig. 12a**, in which we observed gain of loop between two putative enhancer clusters upon CTCF depletion, leading to the upregulation of *Cdiptos* gene. Similarly, we also plotted 37 reproducibly down-regulated DEGs near FINs supported by both studies (**Fig. 4i**). Strikingly, 18 of the recurrent down-regulated DEGs are from protocadherin-γ, a well-known insulator loci (**Fig. 4i,j**). Protocadherin proteins play key roles in the formation of synaptic connections between neurons and the *Pcdh-*γ genes are distinguished by their variable promoters (exon1): γ*a*’s and γ*b*’s are stochastically expressed in individual neurons and the γ*c*’s are constitutively expressed^45, 46, 47,48^. In mESCs, the promoters of γ*a*’s and γ*b*’s are occupied by CTCF, H3K4me3, and the repressive H3K27me3 mark (**Fig. 4j**). A cluster of putative enhancers marked by H3K27ac is present at the 3’ of γ*c* promoters (**Fig. 4j**, highlighted in green) forming loop interactions with the promoters of γ*a*’s and γ*b*’s (**Fig. 4j**, blue arches), which are restricted by a FIN with multiple CTCF peaks (**Fig. 4j**, highlighted in pink). Upon CTCF depletion, the H3K27ac-interactions are weakened, and the *Pcdh-*γ promoter loops extend into a distal H3K27ac-poor region marked by H3K27me3 (**Fig. 4j**), consistent with the downregulation of γ*a* and γ*b* genes. To verify the function of the putative FIN, we designed a pair of sgRNAs to delete the four CTCF peaks in the FIN and confirm the down-regulation of all γ*a* and γ*b* genes (**Fig. 4k-l**). Taking together, these results support the transcription consequences of FINs and nominate “direct targets” of FINs in mESCs.

### Validate FINs with multiplexed CTCF displacement assay

To validate the *cis*-regulatory functions of FINs, we need to verify if the gained loops are truly due to the loss of insulation. Because a FIN often involves multiple CTCF binding sites that contribute to the insulating function collectively, we need to simultaneously perturb multiple CTCF sites in a relatively big region. To achieve this, we combine a multi-sgRNA delivering system name CARGO^49^ with a dCas9-based CTCF-displacement method^47, 50^ to perturb multiple CTCF binding sites in the same cells. In this approach, 4-10 sgRNAs exactly overlapping the cognate motifs of multiple CTCF binding sites are expressed from the same CARGO plasmid. When co-transfected with a dCas9 construct, the sgRNAs can displace CTCF from genome at multiple target sites simultaneously (**Fig. 5a**).

**Figure 5.**
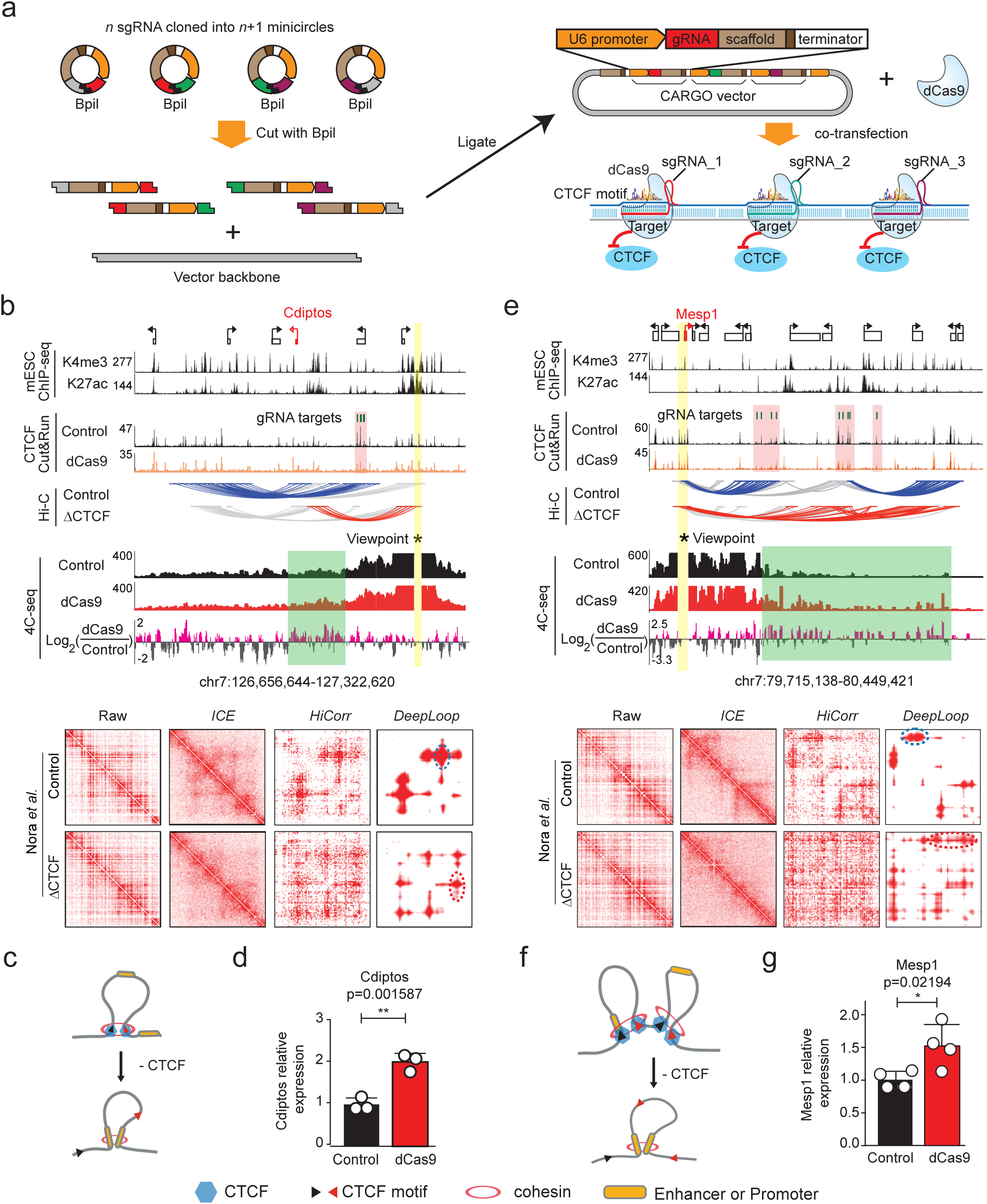
Validate FINs with multiplexed CTCF displacement assay. **a,** Scheme of the CARGO vector that can express multiple sgRNAs. In this system, the sequences of *n* sgRNAs are broken into left and right halves and inserted into (*n*+1) minicircles. The *i*-th minicircle contains the right half of the (*i-1*)-th sgRNA and the left half of the *i*-th sgRNA sequences divided by a *BpiI* restriction site. Final CARGO vector can be assembled by ligating the re-linearized minicircles and vector backbone; each sgRNA is driven by a U6 promoter. Co-transfect CARGO vector and a dCas9-expressing plasmid can displace multiple CTCF binding sites in the same cell. **b,** Validation of a FIN at the *Cdiptos* locus. From the top, 1^st^ and 2^nd^ tracks are H3K4me3 and H3K27ac ChIP-seq data showing the promoter and enhancers in mESC. 3^rd^ and 4^th^ tracks show CTCF ChIP-seq data before and after targeting 4 CTCF motifs in FIN (highlighted in pink) with CARGO. 5^th^ and 6^th^ are Hi-C arches (*DeepLoop* pixels with data from Nora *et al.*) before and after CTCF-depletion (blue: lost loops; red: gained loops). The bottom 3 tracks show 4C-seq data (bait highlighted in yellow) before and after targeting CTCF sites; regions highlighted in green gain 4C signal. The raw and ICE, HiCorr, or DeepLoop processed contact heatmaps are also included. **c,** A model cartoon showing that the gained loop at this FIN locus only involves one CTCF loop. **d,** RT-qPCR results showing that *Cdiptos* is upregulated after CTCF displacement. ** p<0.01; one-side t-test. **e-g,** Validation of a FIN at *Mesp1* locus similar as shown in **b-d**. Note the gained loop at this FIN locus involves at least two CTCF loops (**f**).

We used this approach to validate the gain-of-loop events at two FIN loci. The first FIN is close to *Cdiptos*, a recurrent upregulated DEG (**Fig. 5b**). This example belongs to a simpler type of FIN in which the newly gained loop (**Fig. 5b**, red arches) is blocked by only one FIN (**Fig. 5c**). We used one CARGO construct targeting 4 CTCF motifs and confirmed with CUT&RUN that we successfully weakened the target CTCF peaks but did not affect other CTCF peaks in this region (**Fig. 5b**, highlighted in pink). We performed 4C-seq and confirmed the gain of long-range chromatin interactions consistent with Hi-C data (**Fig. 5b**, highlighted in green). We also validated with RT-qPCR that *Cdiptos* is upregulated upon displacement of target CTCF binding sites (**Fig. 5d**). The second example is more complex as it may involve more than two FINs (**Fig. 5e,f**). In this example *Mesp1* lost its loop interaction to a nearby FIN (**Fig. 5e**, blue arches) and upon CTCF depletion, gained promiscuous distal interactions to a large region that is ∼200kb in size (**Fig. 5e**, red arches). For this locus we used one CARGO construct to target 9 CTCF motifs and successfully weakened the target CTCF peaks (**Fig. 5e**, highlighted in pink). Again, using 4C-seq we confirmed the gain of interactions to the large distal region containing several clusters of putative enhancers (**Fig. 5e**, highlighted in green), and confirmed the upregulation of *Mesp1* with RT-qPCR (**Fig. 5g**). Taking together, these experiments verified the insulating functions of FINs identified from CTCF-degron Hi-C data.

## Discussion

In this study we propose to define functional insulators, or FINs, as CTCF binding sites that actively block the loop formation between flanking *cis*-regulatory elements (**Fig. 3e**). Mapping FINs requires robust detection of newly gained E-P loops, which is challenging for Hi-C analysis because E-P loops are usually weak and hard to detect. With a meta-analysis of three independent CTCF-degron Hi-C datasets, we demonstrate that the subtle gain of loop signal detected by *DeepLoop* are quantitatively robust across different Hi-C or micro-C datasets, therefore proves the feasibility to map FINs reliably with Hi-C at intermediate read depth.

The concept of FIN is distinct from the common assumption that TAD boundaries are insulators. Most TADs disappear upon CTCF depletion, and we typically do not observe gain of new loops across TAD boundaries. In contrast, most of the FINs are anchors of CTCF loops inside TADs in control cells, and we call an CTCF loop anchor “FIN” only if its flanking sequences gain new loops (**Fig. 3e**). Importantly, there are only a few hundred FINs in mESCs, which readily explains why CTCF-depletion only causes mild changes of transcriptome. It should be also noted that the gained E-P loops do not always predict significant gene activation due to complex reasons^51^. Furthermore, target gene expression can be quantitative tuned by multiple enhancers in an additively fashion, and each single enhancer may only have a small effect size^52^.

Nevertheless, we showed that FINs better explain DEGs, especially the recurrent upregulated DEGs in CTCF-depleted cells. This is consistent with the classic definition of insulators as DNA elements that repress enhancer-mediated gene activation.

It remains unclear why only a small subset of CTCF loop anchors are FINs, *i.e.*, why is gain-of-loop only observed at some loci but not others? Our analysis provided some hints. Firstly, the fact that almost all FINs are in compartment A (euchromatin) suggests their dependency on surrounding active chromatin. Notably, promoters and enhancers may play key roles in loop initiation and stalling and are known to co-localize with cohesin and its loading factors^53^. Since genome compartmentalization is a highly dynamic 3D genome feature, we speculate that FINs are also likely to be cell type- or cell state-specific. Secondly, meta-analysis with *Rad21*- and *Wapl*-degron Hi-C data shows that the newly gained E-P loops in CTCF-depleted cells do not form in cohesin-depleted cells and are also likely to be governed by cohesin regulators. It is therefore plausible that local cohesin activity is a key determinant. Finally, *cis-*regulatory elements at the anchors of “gained loops” are enriched with G-rich TF motifs, suggesting that some CREs are particularly capable of forming stable long-range regulatory loops but such function may undergo tight control by insulators. Taking together, all these observations suggest context-specific roles of FINs, and it will be interesting to further explore how FINs exert their specific functions in health and diseases.

## Methods

### Experiments

#### Cell culture

Mouse embryonic stem cells (E14Tg2a) were cultured in DMEM+Glutamax (ThermoFisher, #10566-016) with the addition of 15% Fetal Bovine Serum (ThermoFisher, # SH30071.03), 550µM β-mercaptoethanol (ThermoFisher, #21985-023), 1mM Sodium Pyruvate (ThermoFisher, #11360-070), 1X non-essential amino acids (ThermoFisher, #11140-50), and 10^4^ U of Leukemia inhibitory factor (Millipore, #ESG1107). The cells were maintained at a density of 0.2-1.5x10^5^ cells/cm^2^ and were passaged every 24-48 hours using TrypLE (ThermoFisher, #12563011) on dishes coated with 0.1% gelatin (Millipore, # ES-006-B) at 37°C and 5% CO2. The medium was refreshed daily when cells were not being passaged. Regular checks for mycoplasma infection were performed every 3-4 months, with negative results.

### 4C-seq

4C-seq was conducted following a published protocol^54^. Briefly, 5 million cells fixed with 2% formaldehyde, then quenched with 125 mM glycine. The fixed cells were lysed using a cell lysis buffer containing 50 mM Tris-Cl pH 7.5, 150 mM NaCl, 5 mM EDTA, 0.5% NP-40, 1% Triton X-100, and 1x protease inhibitor cocktail (Roche, Cat#11873580001) for 20–30 minutes on ice.

Following lysis, nuclei were collected by centrifugation at 2,500g for 5 minutes at 4°C and washed once with 1x restriction enzyme buffer. The nuclei pellets were resuspended in 1x restriction enzyme buffer and treated with 0.3% SDS for 1 hour at 37°C with shaking, followed by an additional hour with 2.5% Triton X-100. Chromatin was then digested overnight with the appropriate restriction enzyme at the recommended temperature while rotating in an air bath. The specific restriction enzymes used for each locus are detailed in **Supplemental Table 6**. After digestion, the enzymes were inactivated by heating at 65°C, and nuclei were ligated with 50 μl of T4 DNA ligase (Invitrogen, Cat# 15224–090) in a 7 ml ligation solution at 16°C overnight. Reverse cross-linking was performed by treating the samples with proteinase K to isolate the proximity-ligated DNA. The purified DNA was quantified and subjected to secondary restriction enzyme digestion using roughly 1 unit of restriction enzyme per 1 μg of DNA at the recommended temperature overnight. After enzyme inactivation, the samples were self-ligated with T4 DNA ligase. The ligated DNA was recovered using sodium acetate and ethanol and quantified using the Qubit dsDNA HS assay kit (Thermo Fisher, Cat# Q32851). The 4C templates were amplified using designed primers to generate sequencing libraries. The primer system was modified to be compatible with the Illumina Nextera system, using two sequential PCRs. Locus-specific inverse PCR primers are listed in **Supplemental Table 6**. For each locus, the 4C templates were amplified with locus-specific primers using 200 ng of template per reaction, and products from five parallel amplifications were pooled to generate the final 4C library. A 50 μl PCR product aliquot was purified with homemade Sera-Mag beads. One-fifth of the purified DNA was used for a second PCR with primers N7xx and N5xx, identical to Illumina Nextera sample preparation primers. The final products were purified and sequenced. Reads from the first cutting site were used for data analysis.

### Plasmid cloning

The dCas9 expression vectors used in this study were created using the Cas9 expression vector backbone from the pX330 plasmid (Addgene; plasmid 42230) through the In-Fusion cloning technique. The dCas9 genes were amplified from the pHAGE EF1α dCas9-KRAB plasmid (Addgene; plasmid 50919) via PCR and independently cloned into the AgeI and EcoRI sites of the pX330 plasmid, substituting the Cas9 ORF. Detailed information regarding the primers can be found in **Supplementary Table 5**. All sgRNAs in this study were designed using the CCTop-CRISPR/Cas9 target online predictor.

The Cas9 and *Pcdhg* loop anchor target gRNA expression vector (pDonor_Rosa26_Cas9) was constructed using the backbone of the plasmid pDonor MCS Rosa26 (Addgene, plasmid #37200). An EF-1α-Cas9-P2A fragment, amplified from LentiV_Cas9_Puro (Addgene, plasmid #108100), and a Puromycin-SV40 poly(A) signal fragment, derived from pR26 (Addgene, plasmid #127372), were inserted into the backbone. A paired *Pcdhg* loop anchor target gRNA expression cassette (sequences listed in **Supplementary Table 5**) was then introduced into the construct.

### CARGO assembly

The CARGO array assembly follows a published protocol^49^. The protocol consists of two main steps: constant region preparation and array assembly. In the constant region preparation step, the constant region with proper sticky ends is released by digesting the pGEMT-hU6-spSL (provided by Dr. Joanna Wysocka) plasmid with BsmBI. The released constant region is then separated from the pGEMT backbone on a 1% agarose gel and gel purified using a commercial kit (QIAGEN, Cat#28704). Oligonucleotides (50-60 bp) with variable regions were ordered through IDT (**Supplementary Table 5**). The forward and reverse oligos are annealed and phosphorylated and then stored at -20°C.

In the array assembly step, the annealed chimeric oligo is mixed with the digested constant region, and a minimum of 4-hour ligation with high-concentration T4 ligase (Thermofisher Scientific, Cat#EL0013) is performed. The ligation step yields ligation products, referred to as minicircles. After the stoichiometry check, individual minicircles are treated with plasmid-safe exonuclease to clean up the un-ligated products. Then, the plasmid-safe exonuclease treated individual minicircles are pooled together and purified. The pooled circle mixes are assembled directly into the digested destination vector pX332-mCherry (provided by Dr. Joanna Wysocka) through an optimized golden gate assembly reaction. The product of the golden-gate reaction is then subjected to plasmid-safe exonuclease (Epicentre, Cat#E3101K) treatment. After that, the exonuclease-treated reaction is then transformed into competent cells (Bioline, Cat#BIO-85027). For each plate, individual colonies are picked for plasmid purification using a miniprep kit (QIAGEN, Cat#27204) and test digestion with Acc65I, KpnI, and XbaI. Plasmids with the correct digestion pattern are then verified by Sanger sequencing

### ChIPmentation

We employed ChIPmentation^55^ to map CTCF in various samples. Briefly, cells and tissues were fixed in 1% formaldehyde at room temperature for 15 minutes, followed by glycine quenching.

To isolate nuclei, we lysed cell with lysis buffer 1 (50 mM HEPES; pH 7.5, 140 mM NaCl, 1 mM EDTA, 10% glycerol, 0.5% Nonidet-40, 0.25% Triton X-100, and a protease inhibitor cocktail) for 10 minutes at 4°C. The nuclei were pelleted at 1,800 g for 10 minutes at 4°C and then resuspended in nuclei lysis buffer (10 mM Tris-Cl pH 8.0, 100 mM NaCl, 1 mM EDTA, 0.5 mM EGTA, 0.1% Na-Deoxycholate, 0.5% N-lauroylsarcosine, and a protease inhibitor cocktail) for sonication. For CTCF pulldown, 100-150 µg of chromatin was used. Dynabeads M-280 (Life Technologies, Sheep Anti-Rabbit IgG, #11204D) were washed three times with 0.5 mg/mL of BSA/PBS on ice, then incubated with the CTCF antibody (Abcam, #ab70303) for at least 2 hours at 4°C. The beads/antibody complexes were then washed with BSA/PBS. Pulldown was performed in binding buffer (1% Triton-X 100, 0.1% Sodium Deoxycholate, and a protease inhibitor cocktail in 1X TE) by mixing the beads/antibody complexes with chromatin. After overnight incubation, the beads/antibody/chromatin complexes were washed with RIPA buffer (50 mM HEPES pH 8.0, 1% NP-40, 0.7% Sodium Deoxycholate, 0.5 M LiCl, 1 mM EDTA, and a protease inhibitor cocktail). The beads complexes were then incubated with homemade Tn5 transposase in tagmentation reaction buffer (10 mM Tris-Cl pH 8.0 and 5 mM MgCl2) for 10 minutes at 37°C. To remove free DNA, the beads were washed twice with 1X TE on ice. The pulldown DNA was recovered by reversing crosslinking overnight, followed by purification with PCR Clean DX beads (Aline Biosciences, #C-1003-50). ChIP-seq libraries were generated by amplifying the pulldown DNA with Illumina Nextera primers. Size selection was then performed with PCR Clean DX beads to select fragments ranging from 100 bp to 1000 bp.

### Pcdhg loop anchor deletion

Mouse embryonic stem cells (E14Tg2a) were transfected with the plasmid pX330-sgR26 (Addgene, plasmid #127376), which expresses mammalian codon-optimized Cas9 and the Rosa26 sgRNA, together with the pDonor_Rosa26_Cas9 plasmid, using Lipofectamine™ 3000 reagent (Invitrogen) according to the manufacturer’s instructions. Two days after transfection, the cells were treated with puromycin (2 µg/mL) for three days. Drug-resistant clones were then dissociated into single-cell suspensions, mixed thoroughly, and replated at low density to allow single-colony formation, while maintaining puromycin selection (2 µg/mL). After four days, individual colonies were picked for genotyping using primers specific for the *Pcdhg* loop anchor deletion (**Supplementary Table 5**). Knockout cell clones were identified and selected for subsequent experiments.

### Data analysis

#### Hi-C, micro-C, and 4C-seq

Hi-C and micro-C sequencing reads were mapped to mm10 reference genome separately using bowtie^56^ (v.1.1.2) as previously described^28^. After removing non-unique reads and PCR duplication reads, the bam files are used as input into *HiCorr* in ‘*Bam-process-HindIII*’, ‘*Bam-process-DpnII*’, and ‘*Bam-process-microC*’ modes for bias correction with conventional Hi-C, *in-situ* Hi-C, and micro-C, respectively. After bias correction, we enhance the Hi-C/micro-C data with *DeepLoop*. The *DeepLoop* package provides multiple models optimized for Hi-C data with different read depths^28^. (Note that we use the number of *cis*-contact within 2Mbp as the measurement of read depth.) To choose the appropriate model, we usually selected the model with the closest read depth. The models chosen for each dataset are listed in **Supplementary Table 2**. Because each dataset has a pair of Hi-C experiments (control and KO conditions), if the read depths of the two Hi-C experiments are significantly different, we will down-sample the high-depth dataset to lower depth to reduce potential biases (**Supplementary Table 2**). 4C-seq data were analyzed using pipe4C^54^ (v.1.1.3) to generate bam and bigwig files for data visualization on UCSC genome browser.

### ChIP–seq and CUT&RUN

ChIP-seq and CUT&RUN data were mapped to mm10 (mouse) using bowtie. After mapping step, we remove PCR duplication reads and use Macs2^57^ to call peak and generated bigwig to for data visualization on UCSC genome browser.

### Motif analysis

Motif enrichment analysis was performed by comparing motif occurrences at ATAC peaks located on gained loop anchors against all ATAC peaks. Background motif frequencies were obtained by scanning all ATAC peak sequences using the mouse HOCOMOCO v11^58^ motif database with FIMO (MEME suite)^59^. Motif occurrences in ATAC peaks on gained loop anchors were similarly identified by FIMO scanning. Odds ratios were calculated using Fisher’s exact test to quantify relative motif enrichment. To determine whether motifs were significantly enriched or depleted, a binomial test was applied based on the observed verse expected motif frequencies, and the resulting *p*-values were adjusted to q-values for motif testing correction.

### RNA-seq and mNET-seq

RNA-seq data are aligned to mm10 using HISAT2^60^ and read counts for each gene are called with featureCounts^61^. DESeq2^62^ is used to identify differential expressed genes (DEGs). The DEGs are identified using Wald test with the p-value <0.01 and log2FC>1. For mNET-seq analysis, we followed the customized pipeline from the Hsieh *et al.*^25^ RNA-seq data are aligned to mm10 using HISAT2^60^ and read counts for each gene are called with featureCounts^61^.

DESeq2^62^ is used to identify differential expressed genes (DEGs). The DEGs are identified using Wald test with the p-value <0.01 and log2FC>1. For NET-seq analysis, we followed the customized pipeline from Hsieh *et al.* with the following steps: (i) Trim the adapter using TrimGalore (v.0.6.7) (https://github.com/FelixKrueger/TrimGalore) to remove adapter sequences ‘AGATCGGAAGAGCACACGTCTGAACTCCAGTCAC’ and ‘GATCGTCGGACTGTAGAACTCTGAAC’ at each side of the reads; (ii) Map trimmed read to mm10 using STAR; (iii) Identify the last nucleotide incorporated by Pol II by using mNET_snr (https://github.com/tomasgomes/mNET_snr) to locate the 3′-nucleotide of the second read and the strand sign of the first read. (iv) Annotate and count the reads of each gene using FeatureCounts^61^; (v) Use DEseq2^62^ to quantify the changes of the mNET-seq signal at the gene body between control and KO cells.

### Pairwise loop strength comparison

For each pair of CTCF/cohesin-degron study, we took top 500k loop pixels from the *DeepLoop* enhanced contact matrices of control and IAA/dTAG-treated samples separately, then union them together. The loop strength of each pixel in the two experiments are normalized linearly:

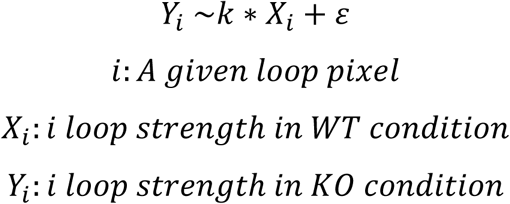

However, because CTCF- and *Rad21*-depletion leads to massive loss of loop signal, much fewer loop pixels are expected in the IAA/dTAG-treated cells than in control cells. To adjust for this, we have to manually pick a regression slope *k* based on the shape of the scatter plots. The slopes we chose are listed in **Supplementary Table 2**. We then use the minimum strength of the control cell loop pixels to adjust the minimum cutoff to call loop pixels in KO samples: 𝑌_min_ = 𝑘 ∗ _min_(𝑋_𝑖_). In another word, a pixel is called as loop pixel in KO sample only if its strength is bigger than 𝑌_min_ This step leads to much fewer loop pixels in CTCF- and *Rad21*-depleted cells. We typically obtain ∼600k chromatin loop for each study. We used a simple fold change cutoff (FC > 4) to define gained or lost loop pixels. Finally, we used the following criteria to call retained loops: (i) the strength of the pixel in both WT and KO conditions must be higher than their minimum cutoffs (𝑋_𝑖_ > 𝑋_min_ & 𝑌_𝑖_ > 𝑌_min_); (ii) the absolute value of residual must be higher than the adjusted minimum of loop pixels in KO samples (|𝜀| < 𝑌min).

### Classify chromatin loops based on epigenome data

For the purpose of being consistent, we intersected loop anchors with the same set of mESC CTCF, H3K27ac, H3K4me3, H3K27me3 and H3K36me3 ChIP-seq peaks from ENCODE^63^ to classify the loop pixels. We classification loop pixels in the following order:

1. EP-only: both anchors have either H3K4me3 or H3K27ac mark, but they do not have CTCF peaks.
2. CTCF-EP: after excluding EP-only, look for pixels which only one anchor has CTCF, and the other anchor has either H3K4me3 or H3K27ac marker.
3. CTCF-only: after excluding EP-only and CTCF-EP, look for pixels with at least one anchor bound by CTCF.
4. H3K27me3: after excluding (1) - (3) above, look for pixels with at least one anchor bound by H3K27me3.
5. H3K36me3: after excluding (1) - (4) above, look for pixels with at least one anchor bound by H3K36me3.
6. The rest pixels are classified as “other”.

### Define Loop-gaining loci, FINs, FIN loci, and FIN-blocked regular enhancers and super enhancers (SE)

Loop-gaining loci are defined from “gained” loop pixels in CTCF-depleted cells. We identified thousands of “gained” loop pixels and each pixel represents a pair of 5kb-bins. However, these pixels often occupy overlapping regions. We therefore merge all overlapping pixels into one region and define them as loop-gaining loci. This step allows us to count the number of regions in the genome containing “gained” loops. In our study, we have ∼300 loop-gaining loci after merging ∼5,000 recurrent “gained” loop pixels.

We define FINs as a subset of CTCF binding sites that satisfy the following criteria: (i) they located at the anchors of CTCF loops that are lost upon CTCF depletion; (ii) the FIN should be inside a newly gained loop in CTCF-depleted cells; (iii) the other anchor of the CTCF loop should be outside of the newly gained loop; (iv) CTCF sites satisfying (i-iii) within 20kb of each other will be merged into one FIN. Note that we do not have limits on the number of CTCF sites to define FINs. Therefore, although a FIN can be just one single CTCF site, most FINs have multiple CTCF sites.

As described above, all FINs must be inside loop-gaining loci. “FIN loci” are “loop-gaining loci” that containing FINs. In fact, almost all the “loop-gaining loci” are also “FIN loci”. We therefore often use the two terms interchangeably.

We call enhancers as H3K27ac peaks and use a published list of super enhancers in mESCs^64^ (**Supplementary Tables 11 & 12**). This allows us to determine the types of enhancers blocked by FINs.

### Recurrent loop-gaining pixels and loop-gaining loci in CTCF depleted cells

A recurrent loop-gaining pixel is called if it satisfies FC > 4 in at least two studies. We define the location of each loop pixel as the entire region between the left and right anchors (including the anchors themselves). To define loop-gaining loci, we merge all gained loop pixels from the CTCF-degron studies using the “merge” option in *bedtools*; each loop-gaining locus usually has multiple loop pixels. Loop loci with gap size < 20kb will be merge into one locus. In this study, we merged all the recurrent loop-gaining pixels to obtain 299 recurrent loop events. To test the consistency of loop-gaining loci between independent CTCF-degron study, we call loop-gaining loci for each individual CTCF-degron study and the share loop-gaining loci from two different studies are called if they have at least one base overlap.

### Test the recurrence of loop-gaining loci from CTCF-degron in Rad21- and Wapl- degron data

Nearly all loop-gaining loci observed in Hi-C degron studies have at least one E-P loop pixels. We further classify the loop-gaining loci depending on the number of CTCF loops separating the two anchors of the gained E-P loops. For all loci, we also manually examined all loci to confirm the classification. Several loci cannot be clearly classified into the categories shown in **Fig. 3d** because they have different marks or have multiple events. To test if a loop gaining locus is recurrent in one of the CTCF/cohesin-degron study in **Fig. 3d**, we examine all loop-gaining pixels in the locus and if one of pixels also gain signal it will be considered recurrent.

### Compartment and TADs in mESCs

Hi-C compartment analysis is performed in WT mESCs following previous publication^65^. After QC step, we constructed 500kb resolution contact matrices for each chromosome and then normalized them using the genome distance. We next generated correlation matrices for each chromosome and performed the principal component analysis on the correlation matrix and then assigned the genome into two compartments by using PC1. We used the density of transcription starting sites (TSS) of each 500-kb bin to determine compartment A and compartment B.

### TAD boundary calling with Insulation score or Directionality index (DI) methods

Insulation scores were computed for contact matrices using 40kb bins and 2Mb sliding window using the *cooltools*^66^ package. Insulating loci were identified through a local minima detection procedure based on peak prominence. Highly insulating regions corresponding to strong TAD boundaries were defined using the Li thresholding method implemented in the *cooltools* insulation score module. In this method, TAD boundaries are a list of 40kb bins.

We also called TAD domains with control mESC Hi-C data using the DI method described by Dixon *et al.*^1^, which quantifies the degree of upstream or downstream interaction bias for each genome bin. Directionality indices were calculated also with 40kb bins using a 2Mb sliding window. A hidden Markov model (HMM) was applied to infer hidden bias states across the genome, classifying each bin as upstream-biased, downstream-biased, or neutral. Consecutive bins sharing the same bias state were merged into domains. The start and end positions of each domain are defined as TAD boundaries. In this way, the TAD boundaries will be also a list of single bases. The TAD boundaries from the WT samples of the three datasets are listed in **Supplementary Table 9**.

To compare the TAD boundaries from two methods, we consider two TAD boundaries overlapping if their distance gap is smaller than 40kb. The same distance cutoff (40kb) is also used to decide if a FIN overlaps TAD boundary.

### Ideogram plot

We generate ideogram plots with Rideogram^67^ to visualized the locations of recurrent gaining-loop pixels, compartment, and TADs.

### Density and enrichment of DEGs upon CTCF-depletion

For any region in the genome, the density of DEGs is defined as the average number of DEGs per Mb-region. We computed DEG density for TAD boundary +/- 100kb, FIN +/- 100kb, gained loop anchors, lost loop anchors, and retained loop anchors. We also compute the enrichment of DEGs at regions of interest as their DEG density divided by the DEG density in the whole genome.

### Hi-C data visualization

Heatmap was used to visualize the raw, ratio, and *DeepLoop*-enhanced heatmaps. We use the following strategies as described before^28^ to determine the color scales these heatmaps. (i) Raw or ICE-normalized heatmaps represent raw read counts; the brightest red color indicates the 98th percentile of the contact matrix. Color is proportionally scaled down to one read (white). (ii) HiCorr heatmap represents the ratio value after Hi-C bias correction; the brightest red color indicates at least twofold enrichment of a given pixel comparing to its background. Color is proportionally scaled down to onefold (no enrichment). (iii) DeepLoop heatmaps outputs ‘transformed fold change’ that represents only relative levels of signal enrichment (for example, a value of one may no longer represents no enrichment). We therefore set the brightest red color as the lower limit of the top 300k pixels genome wide. Color is proportionally scaled down to half of that lower limit or onefold, whichever is higher. All loop arches shown in figures were generated with UCSC Genome Browser^68^.

## Data availability

Accession numbers for third-party data used in this study can be found in **Supplementary Table 1**. The raw data of 4C–seq and ChIP-seq generated in this study, and reanalyzed published data, can be found at accession number GSE243728. The 18 Hi-C and micro-C datasets analyzed by *DeepLoop* can be found at https://hiview.case.edu/public/DeepLoop/.

To review GEO accession GSE243728:

Go to https://www.ncbi.nlm.nih.gov/geo/query/acc.cgi?acc=GSE243728 Enter token uvixkcaajpajpih into the box

## Code availability

The code is available is available at github (https://github.com/JinLabBioinfo/FIN_project).

## Acknowledgements

This work was supported by grants from NIH R01HG009658 and R01CA267872 (to F.J.), R01DK113185 (to Y.L.), R01DK131437, R01HG012384, and UG3NS132061 (to Y.L. and F.J.). F.J. is also supported by a subaward from NIH project (U01AG072579) through University of Miami and partially by several other NIH grants R01CA252224, R01GM148662, and P01CA272161. Y.L. is also partially supported by NIH grants R01CA264320. This work made use of the High-Performance Computing Resource in the Core Facility for Advanced Research Computing at Case Western Reserve University.

## Author Contributions Statement

F.J. and Y.L conceived the project and secured funding for this project. J.C. designed and performed the FIN validation experiments, W.X. and X.L. performed the data analysis. L.L. help with the experiments and S.Z. and X.L. helped with the data analysis. F.J., W.X., J.C.,X.L., and Y.L. wrote the manuscript. F.J. and Y.L. also provided critical feedback and interpretation of results as well as helping to revise the manuscript.

## Supplementary figure legend

**Supplementary Figure 1. Compare the compartments and TADs between orthogonal CTCF-degron datasets from Hi-C, in situ Hi-C, and micro-C.**

**a-c,** Compartment analysis of CTCF-degron Hi-C datasets at 500kb resolution. **a,** Correlation heatmaps of the three CTCF-degron Hi-C datasets at 500kb resolution showing the compartments in control and CTCF-depleted cells. PC1 values are shown on top of each heatmap indicating compartment A (positive PC1) and B (negative PC1). **b**, Scatter plots comparing the PC1 values for each 500kb bin before and after CTCF depletion. **c,** Top row: scatter plots comparing the PC1 values of Hi-C data in control cells from three independent studies; bottom row: scatter plots comparing the PC1 values of Hi-C data in CTCF-depleted cells from three studies. **d-f,** TAD analysis of CTCF-degron Hi-C datasets at 40kb resolution. **d,** ICE normalized contact heatmaps of the three CTCF-degron Hi-C datasets at 40kb resolution. TADs are recognizable as squares along the diagonals. Note the loss of TADs in CTCF-depleted cells. **e,** Scatter plots compare the directionality index (DI) values of all 40kb bins before and after CTCF-depletion in all three studies. Note that the DI values of many bins become close to zero in CTCF-depleted cells, indicating that these bins lost directional biases, consistent with massive loss-of-TAD upon CTCF-depletion. **f,** Top row: scatter plots comparing the DI values of all 40kb bins in control cells from three independent studies; bottom row: scatter plots comparing the DI values of all 40kb bins in CTCF-depleted cells from three studies. Note that although the R-squares between Nora et al. and Kubo et al. are rather high (R-square 0.718 and 0.612 for control and ΔCTCF cells), the DI values from micro-C data (Hsieh et al.) deviate significantly from the other two studies (R-square 0.295∼0.458). In micro-C data, more bins show polarized DI values (*i.e.*, stronger directionality biases), consistent with the general belief that with ICE-based pipeline, micro-C identifies more loops than Hi-C or in situ Hi-C.

**Supplementary Figure 2. Compare the loop-level analyses of orthogonal Hi-C datasets with ICE, HiCorr, and DeepLoop, related to Fig. 1b**.

**Part I.** Cross-platform comparison of CTCF-degron datasets. **a-c,** Loop analysis of three CTCF-degron Hi-C datasets with ICE at 5kb resolution. **a,** ICE heatmaps of the three CTCF-degron Hi-C datasets at 5kb resolution. Note that there are more noises in Hi-C (Nora) and *in situ* Hi-C (Kubo) heatmaps than in micro-C (Hsieh) heatmaps. **b,** Scatter plots of ICE heatmap pixels comparing the control cells contact heatmaps from three independent studies. Note that the consistency between orthogonal Hi-C protocols is low (R-square values close to zero). **c**, Scatter plots comparing the ICE heatmaps of control and ΔCTCF cells in three different studies (5kb resolution). Every dot in the scatter plot represents one pixel in the contact matrix and the x-axis and y-axis indicate the intensity of the pixel in two experiments. Note that ΔCTCF cells have many pixels stronger than WT cells because the read-depth of micro-C data in ΔCTCF cells is higher than in control cells. **d-f,** Similar as **a-c** but the comparisons are between ICE heatmaps after correcting distance biases, which are observed/expected ratio heatmaps. Note that in **f**, the control vs. ΔCTCF scatter plot for Hsieh *et al.* correctly show stronger loop pixels in control cells than ΔCTCF cells. However, the scatter plots in **e** show that the consistency between independent studies remains poor with R-square values still close to zero, indicating that the pixel-level correlations are dominated by random noise pixels. **g-i,** the same comparisons between HiCorr-corrected heatmaps, which are also distance-corrected ratio heatmaps. Note that the scatter plots in **i** show that in each of the control vs. ΔCTCF comparisons, we start to observe retained pixels with similar strength before and after CTCF depletion. Furthermore, the scatter plots in **h** also show that HiCorr achieves higher R-square values across different Hi-C platforms. **j-l,** comparisons between *DeepLoop* heatmaps. Note that DeepLoop highlights loop pixels and suppresses noise pixels (**j**). The retained pixels and weakened pixels are more clearly recognizable in each of the control vs. ΔCTCF scatter plots (**l**). DeepLoop also shows the best cross-platform consistency at pixel level (**k**).

**Part II.** Cross-platform comparison of Rad21-degron datasets. Same as Part I and all conclusions are the same.

**Part III.** Cross-platform comparison of Wapl-degron datasets. Same as Part I and all conclusions are the same.

**Supplementary Figure 3. Correlation matrices comparing the profiles of dynamic loops from all nine pairs of degron experiments, related to Fig. 1c**.

For each degron experiment, we first obtain a pair of 5kb resolution Hi-C contact matrices in control and depletion cells. Next, we computed the delta matrices (depletion – control) for all nine experiments. A correlation matrix is then generated between the nine delta matrices. **a-d** are correlation matrices when contact matrices are generated with different pipelines.

**Supplementary Figure 4**. **Recurrent gain of E-P loop at *Notch1* locus between independent CTCF-degron Hi-C studies, related to Fig. 1d,e**.

**a,** Contact heatmaps of the CTCF-degron Hi-C data by Kubo *et al.* at *Notch1* locus with *ICE*, *HiCorr* and *DeepLoop*. Red dashed circle highlights the same locations where we observed a gained E-P loop in the contact heatmap from Nora *et al.* (shown in **Fig. 1d**). **b**, Genome browser tracks showing ChIP-seq data and Hi-C loops from Kubo *et al.* Blue highlighted region indicates *Notch1* gene. Pink region highlights the potential insulator region that blocks the formation of *Notch1* E-P loop. **c,** RNA-seq results from Kubo *et al.* are also shown as bar plot. p-value: one-side Wald test provided by DE-seq2. **d-f,** the same analysis as **a-c** at *Notch1* locus using data from Hsieh *et al.* Note that Hsieh *et al.* used mNET-seq for transcriptome profiling. **g,** Contact heatmaps of *Notch1* locus from two *Rad21*-degron datasets (Kriz *et al.* and Hsieh *et al.*) in addition to the one included in **Fig. 1d** (Rhodes *et al.*). **h,** Contact heatmaps of *Notch1* locus from two *Wapl*-degron datasets (Kriz *et al.* and Hsieh *et al.*) in addition to the one included in **Fig. 1d** (Liu *et al.*).

**Supplementary Figure 5. Meta-analysis of independent CTCF-degron Hi-C datasets to examine the consistency and epigenetic features of dynamic loops.**

**a**, Using the *in situ* Hi-C data from the CTCF-degron study by Kubo *et al.*, left: scatterplot comparing the strength of loop pixels between WT and CTCF-depleted mESCs; middle: the same scatterplot as left but highlight the lost (blue), retained (black), and gained (red) loop pixels; right: violin plot comparing the size of loop pixels. **b,** Pie charts showing the components of lost, retained and gained loop pixels based on epigenetic marks. **c**, Scatter plots in the left column highlight the lost, retained, and gained loop pixels from Kubo *et al*., the right two columns show the quantitative changes of the corresponding pixels in Nora *et al.* or Hsieh *et al*. **d-f,** The same analyses as **a-c** are performed with CTCF-degron micro-C data by Hsieh *et al*.

**Supplementary Figure 6. Examples of chromatin loop classification.**

Each panel shows the genome browser tracks with loop arches. These examples include **a**, EP loop; **b**, CTCF-EP loop; **c**, CTCF loop; **d**, H3K27me3 loop; **e**, H3K36me3 loop.

**Supplementary Figure 7. Meta-analysis of independent *Rad21*- and *Wapl*- degron Hi-C datasets to examine the consistency and epigenetic features of dynamic loops.**

**Part I:** Meta-analysis of *Rad21*-degron datasets. **a,** Left and middle panels define the lost, retained, and gained loop pixels upon *Rad21*-depletion from the study by Rhodes *et al.*, and the right panel compares their loop size distribution. **b,** Pie charts showing the components of lost, retained and gained loops based on epigenetic marks. **c,** Scatter plots testing if the lost, retained and gained loop pixels from Rhodes *et al.* are quantitatively consistent in the other two *Rad21*-degron studies. **d-f,** the same analyses as **a-c** are performed with *Rad21*-degron Hi-C data by Kriz *et al.* **g-i,** the same analyses as **a-c** are performed with *Rad21*-degron micro-C data by Hsieh *et al*.

**Part II:** Meta-analysis of *Wapl*-degron datasets. **j-r,** Same as Part I except that the analyses were done with Wapl-degron datasets. Note there is an overall trend that *Wapl*-depletion causes more “gained” loops.

**Supplementary Figure 8. “Gained” long-range H3K27me3 loops in Rad21-depleted cells from Rhodes *et al.* exist before Rad21-depletion in two other studies.**

Two examples are shown in **a** and **b**. In each example, Hi-C data are shown as both contact heatmaps (DeepLoop enhanced) and loop arches. Browser tracks of CTCF, H3K27ac, H3K4me3, and H3K27me3 are also included. Note that the gained H3K27me3 loop in Rhodes et al. already exist before Rad21-depletion in two other studies.

**Supplementary Figure 9. Meta-analysis reveals the fate of dynamic loops from CTCF-degron studies upon depletion of cohesin factors.**

**a,** The lost, retained, and gained loop pixels from one CTCF-degron Hi-C data (Kubo *et al.*) are plotted onto Hi-C scatter plots from *Rad21*- or *Wapl*-degron studies to examine their quantitative changes. **b,** Same analyses are done with the lost, retained, and gained loop pixels from the CTCF-degron Hi-C data by Hsieh *et al*.

**Supplementary Figure 10. Examples of loop gaining loci, related to Fig. 3d**.

**a,** An example of Type I EP loop gaining loci, in which one of the two E/P sites is insulated in a CTCF loop while the other site is outside the CTCF loop; **b,** An example of Type II EP-loop gaining loci, in which two E/P sites are insulated in two CTCF loops; **c,** An example of Type III EP loop gaining loci, in which neither of the two E/P sites are insulated by CTCF loops; **d,** An example of CTCF loop gaining loci, in which the newly gained loop are between incompletely degraded CTCF sites.

**Supplementary Figure 11. Limited overlap between FINs and TAD boundaries, related to Fig. 3f,h,i.**

**a,** Venn diagrams comparing the TAD boundaries called by two methods from three studies. **b,** Pie chart showing the percentage of FINs that overlap TAD boundaries called from different datasets and algorithms. **c,** Compare insulation scores at FINs and TAD boundaries in different studies.

**Supplementary Figure 12. Example contact heatmaps at putative insulator target genes, related to Fig. 4h,j**.

**a,** Raw, *ICE*, *HiCorr*, and *DeepLoop* contact heatmaps at *Cdiptos* locus in control and CTCF-depleted cells from Hsieh *et al.* and Nora *et al.* **b,** Similar as **a** but the heatmaps are at the Pcdh-γ locus.

