## Supplementary figures and images for "Cross-platform Hi-C meta-analysis identifies functional insulators that actively block enhancer-promoter interactions"

### Supplemental Figure 1

Supplementary Figure 1

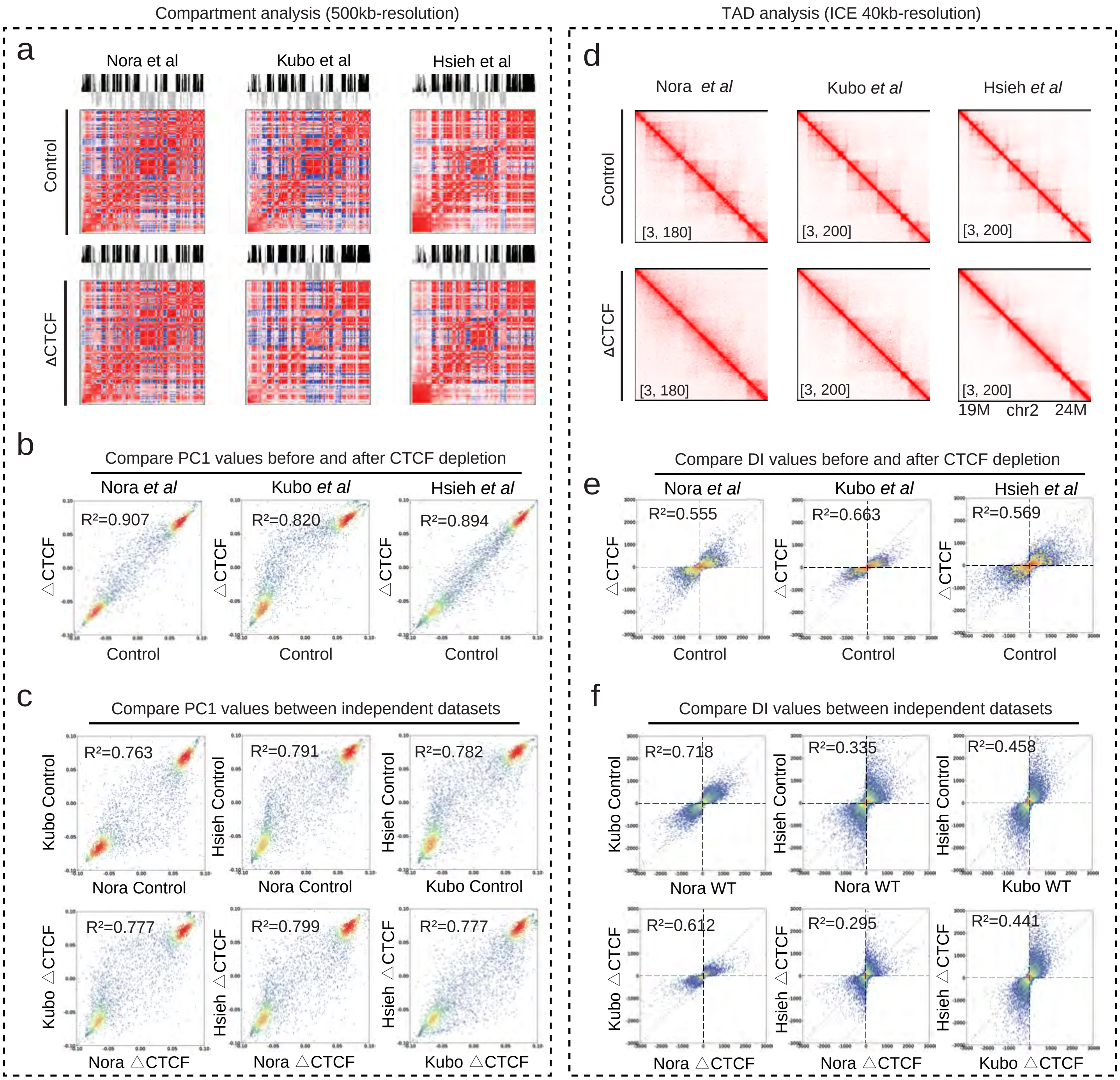

### Supplemental Figure 2

#### Supplementary Figure 2

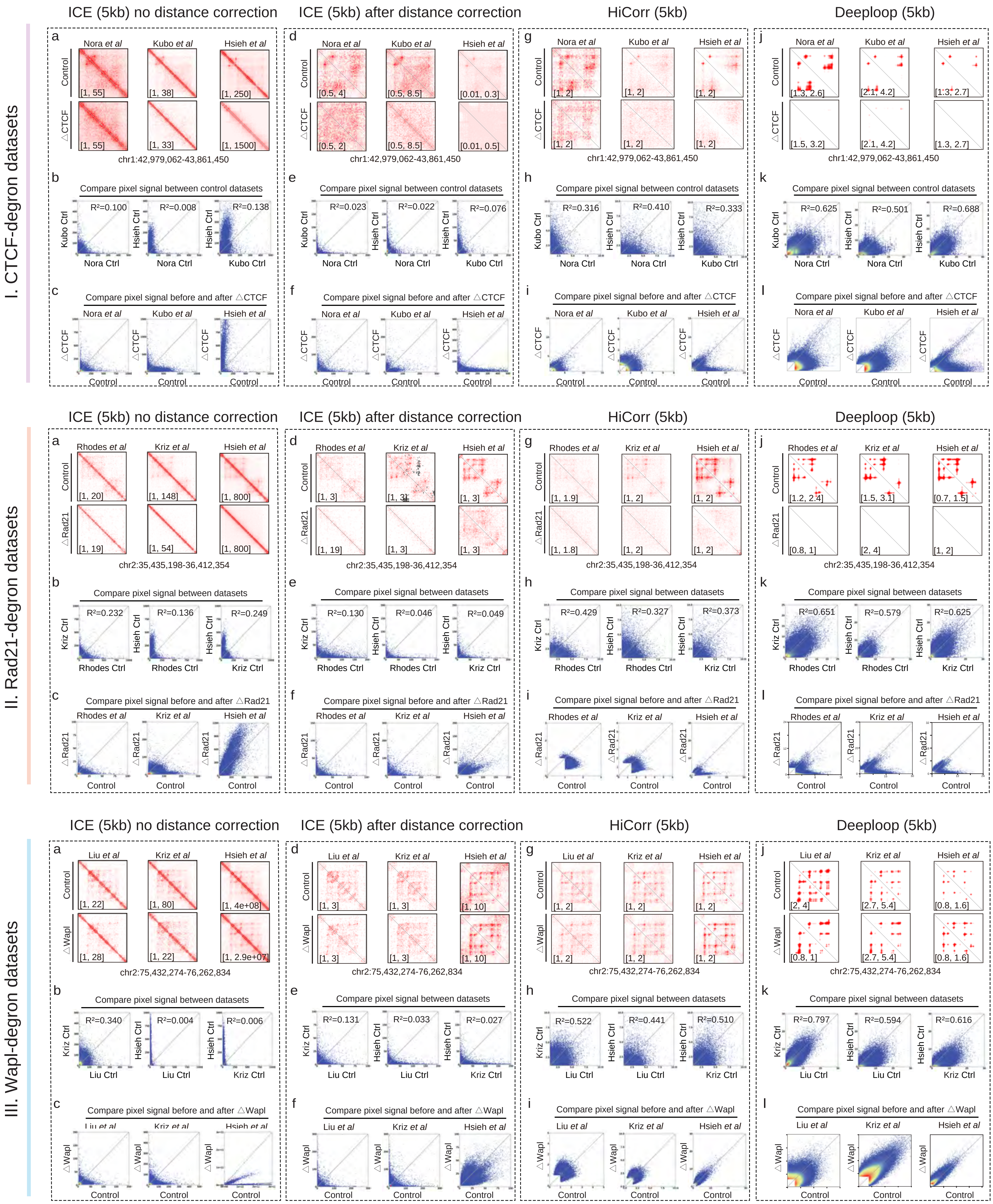

### Supplemental Figure 3

#### Supplementary Figure 3

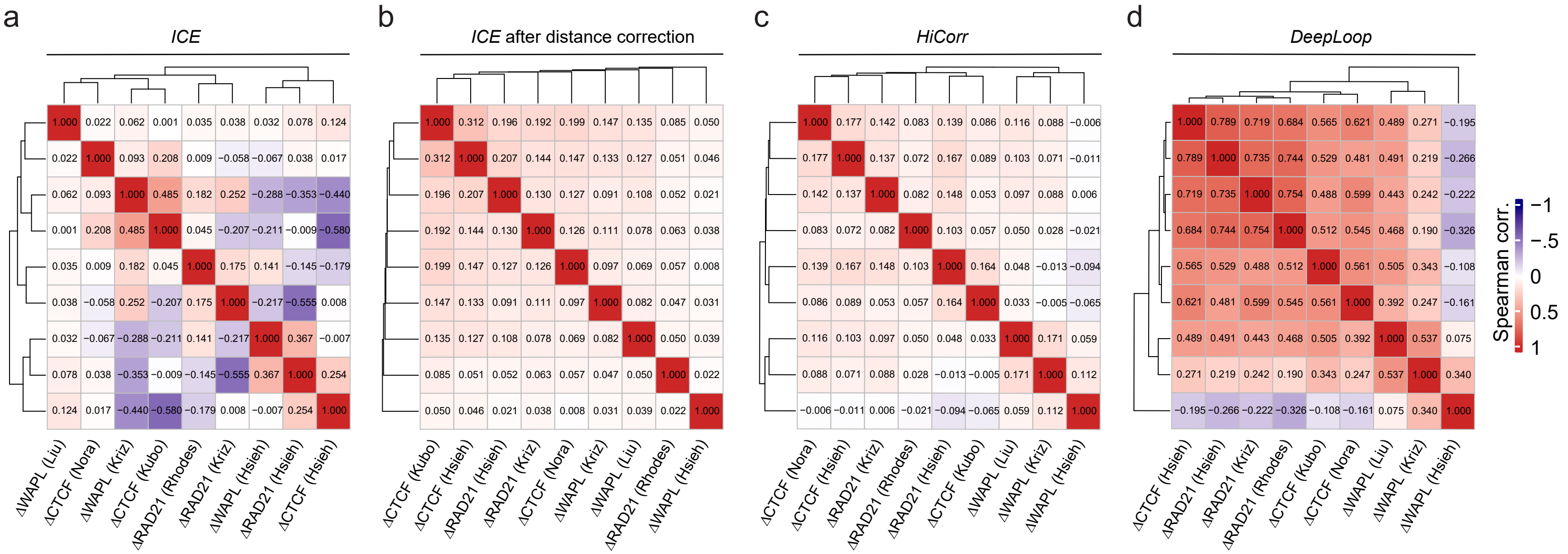

### Supplemental Figure 4

#### Supplementary Figure 4

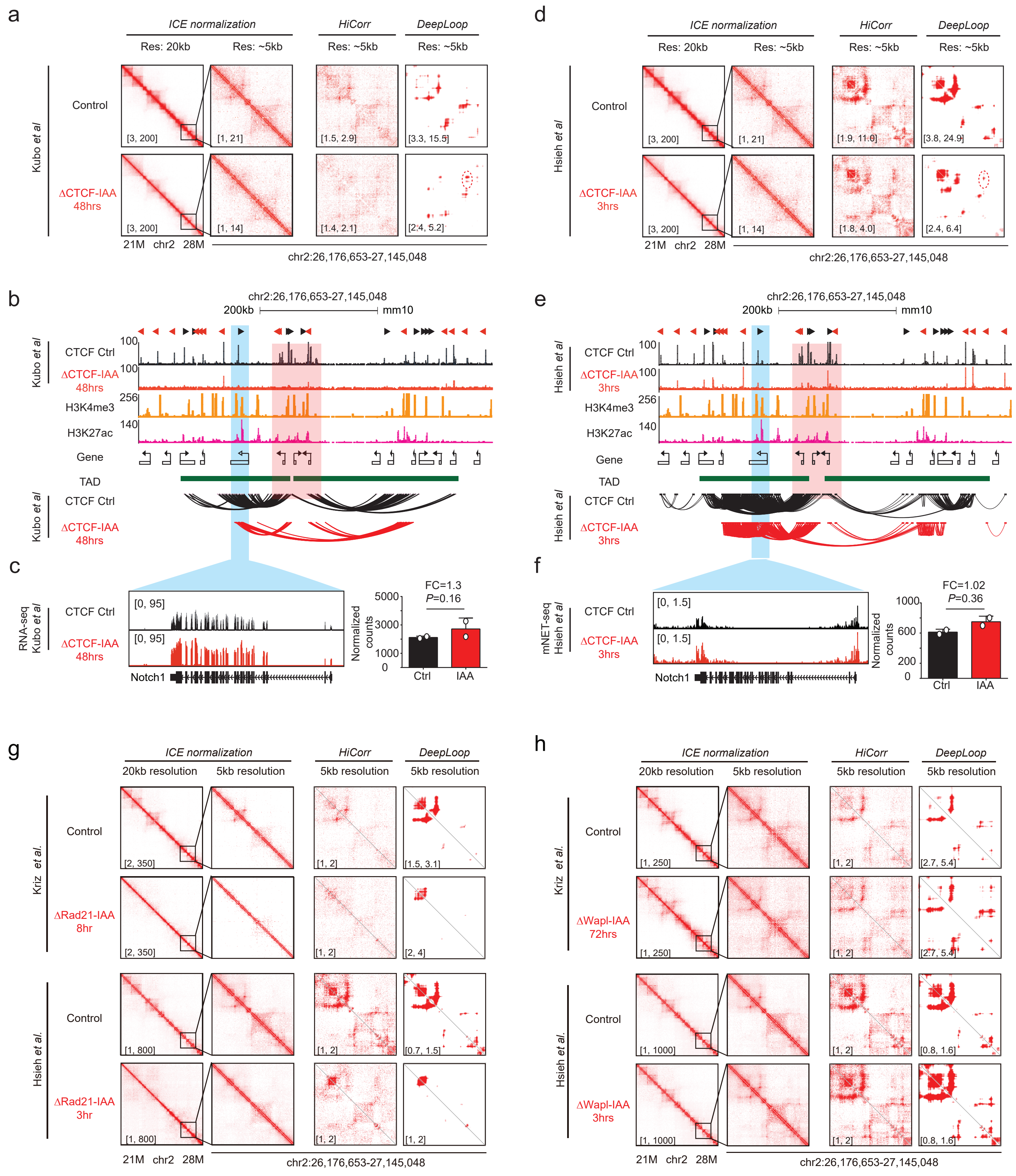

### Supplemental Figure 5

##### Supplementary Figure 5

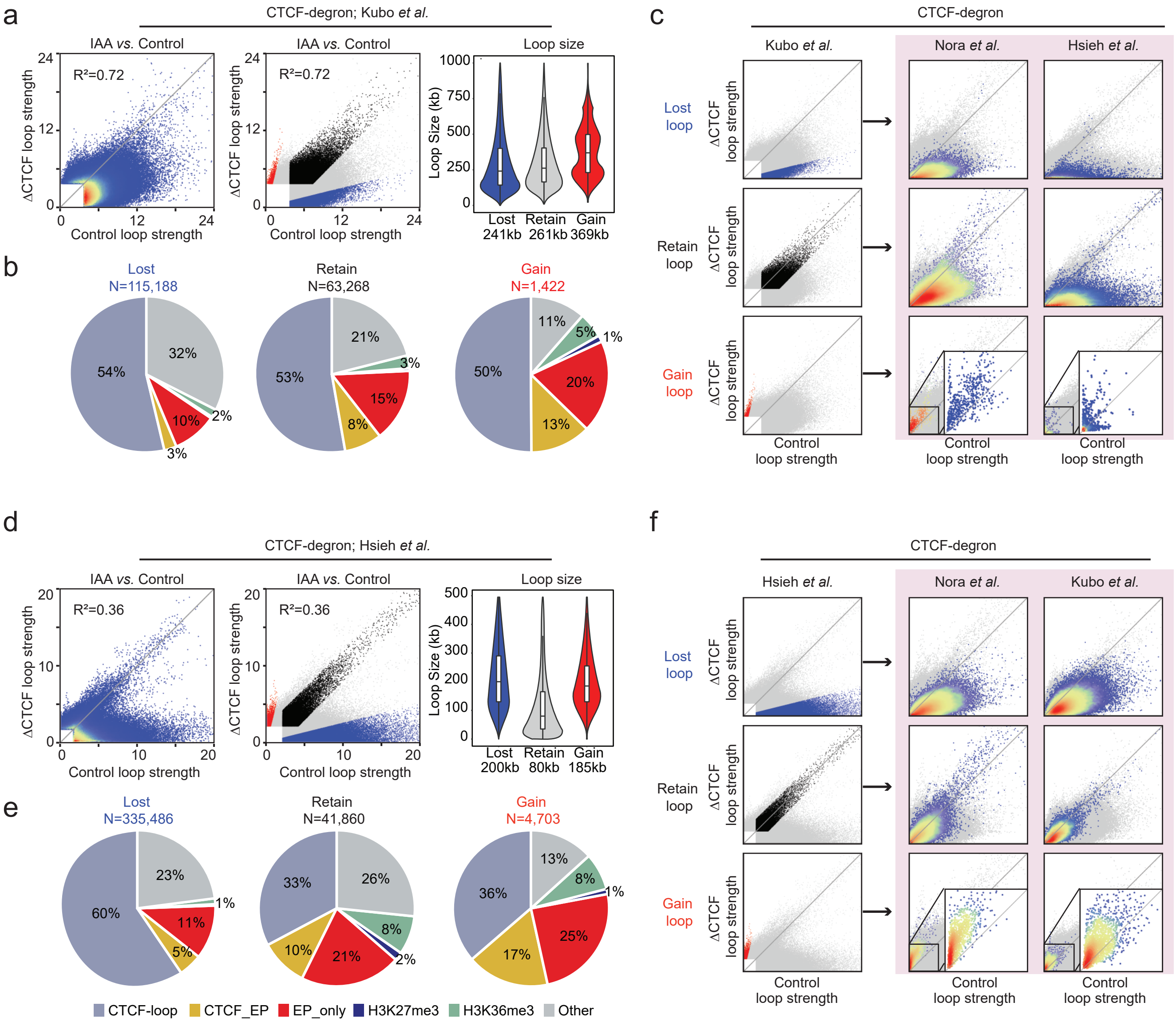

### Supplemental Figure 6

#### Supplementary Figure 6

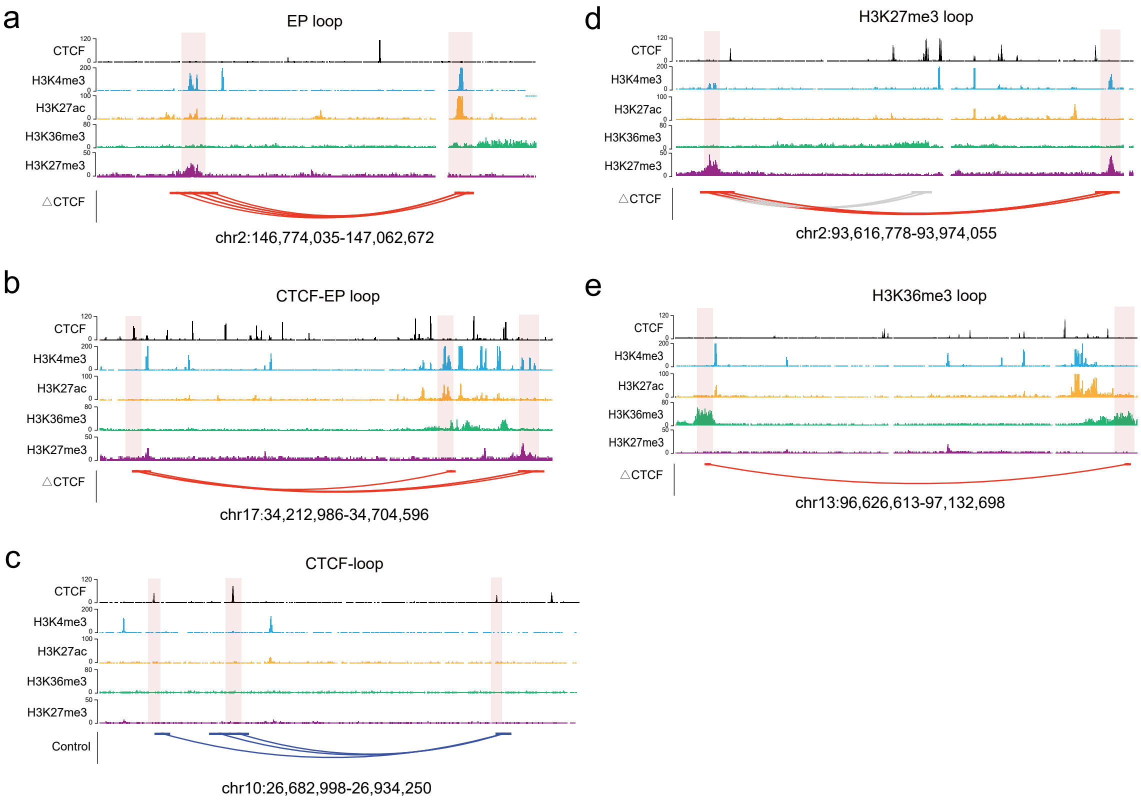

### Supplemental Figure 7

#### Supplementary Figure 7

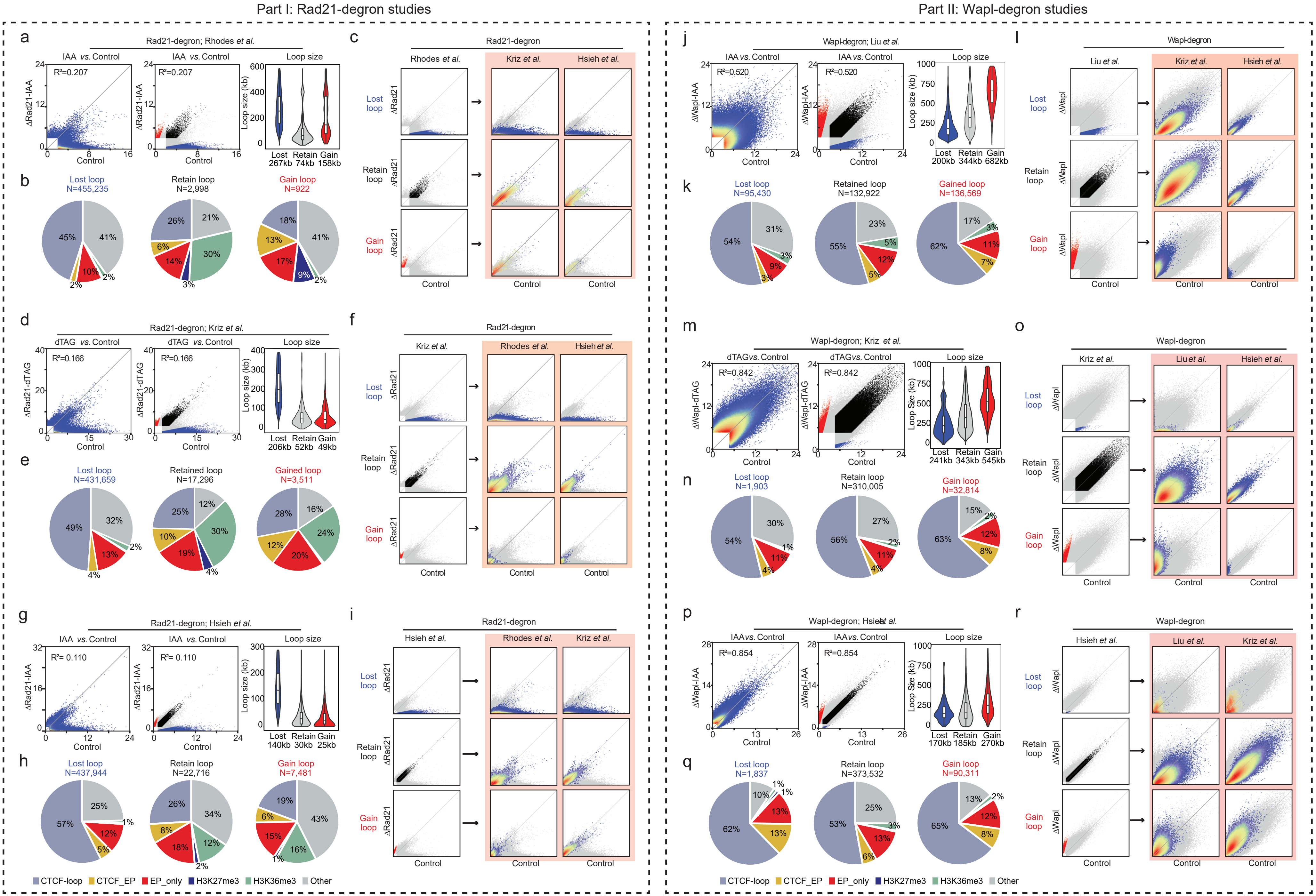

### Supplemental Figure 8

Supplementary Figure 8

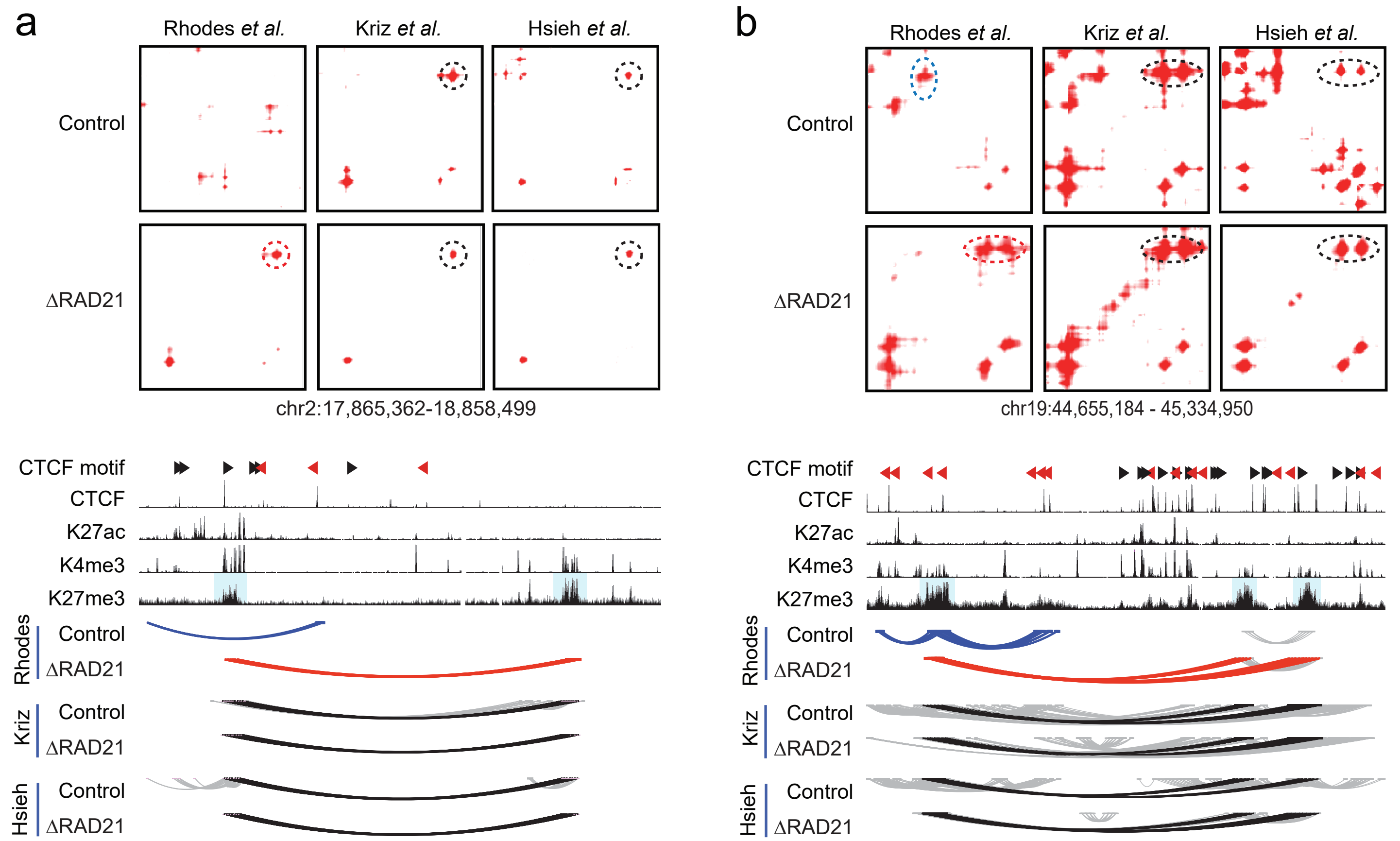

### Supplemental Figure 9

#### Supplementary Figure 9

a

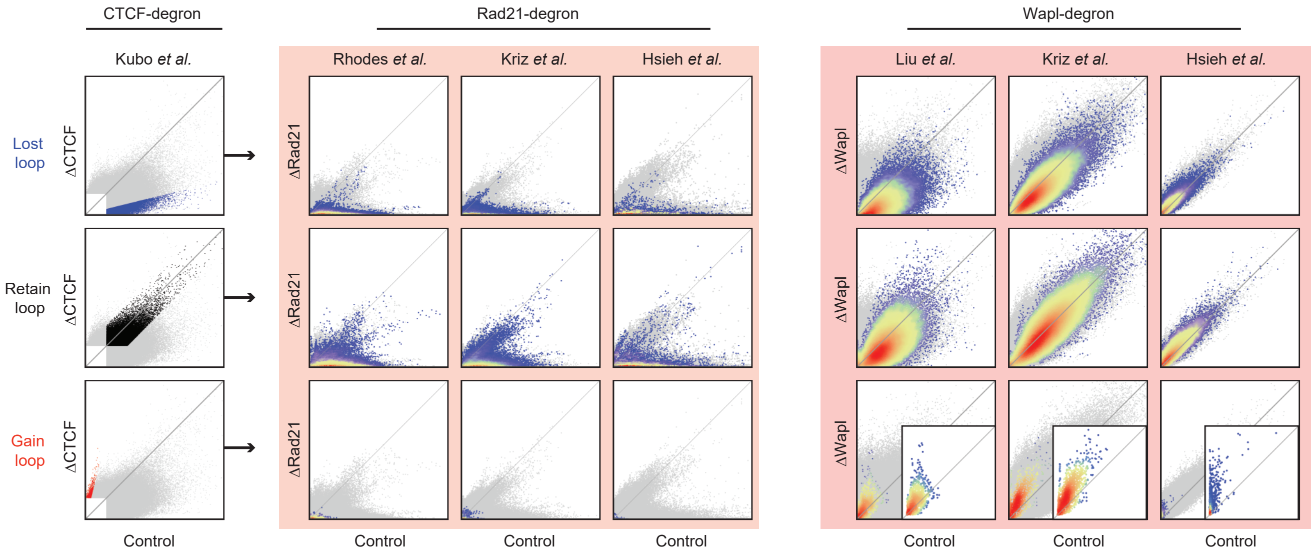


b

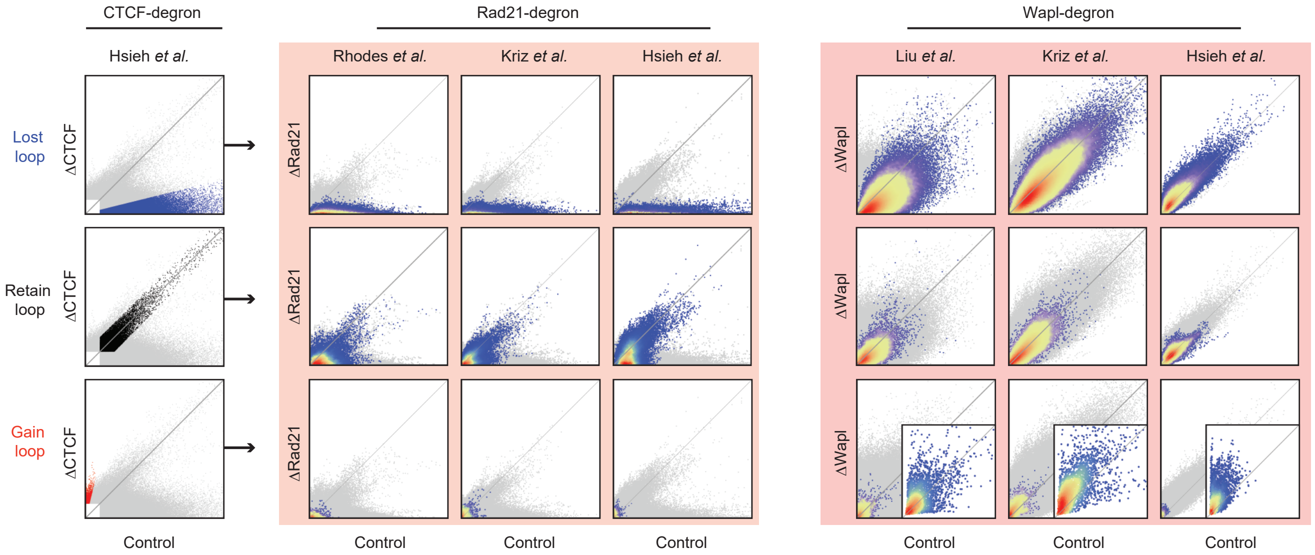

### Supplemental Figure 10

#### Supplementary Figure 10

a

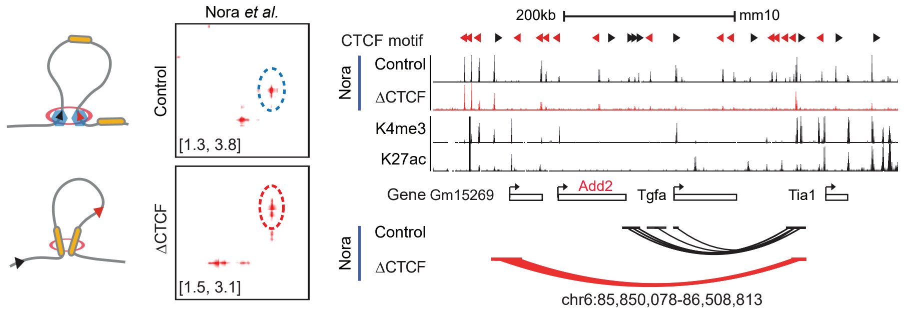


b

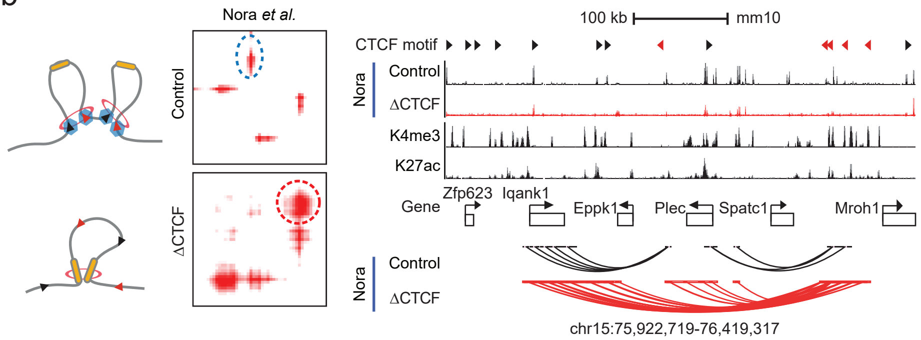


c

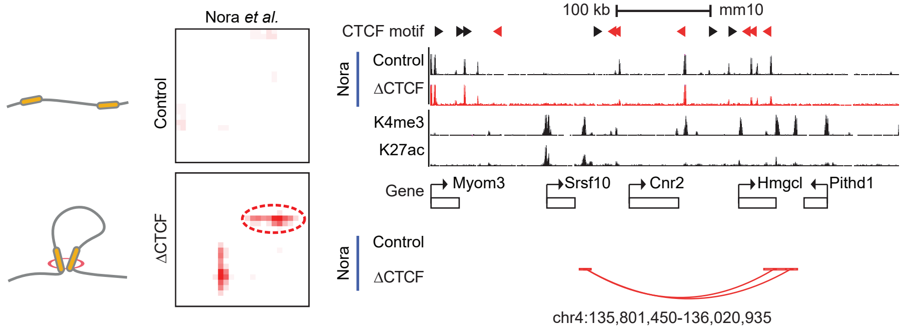


d

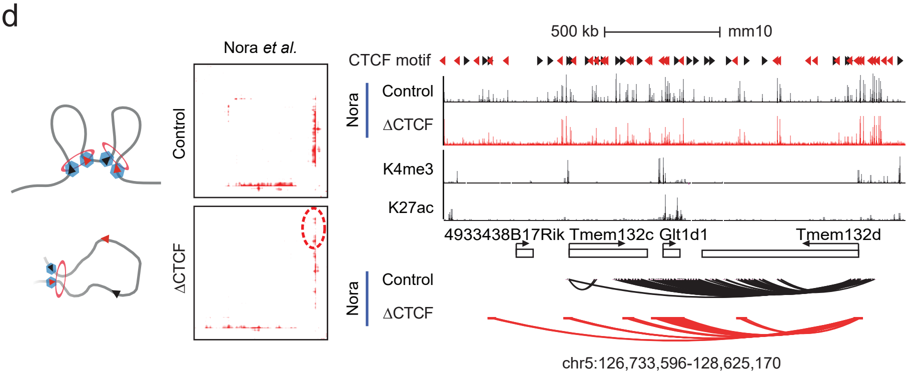

### Supplemental Figure 11

#### Supplementary Figure 11

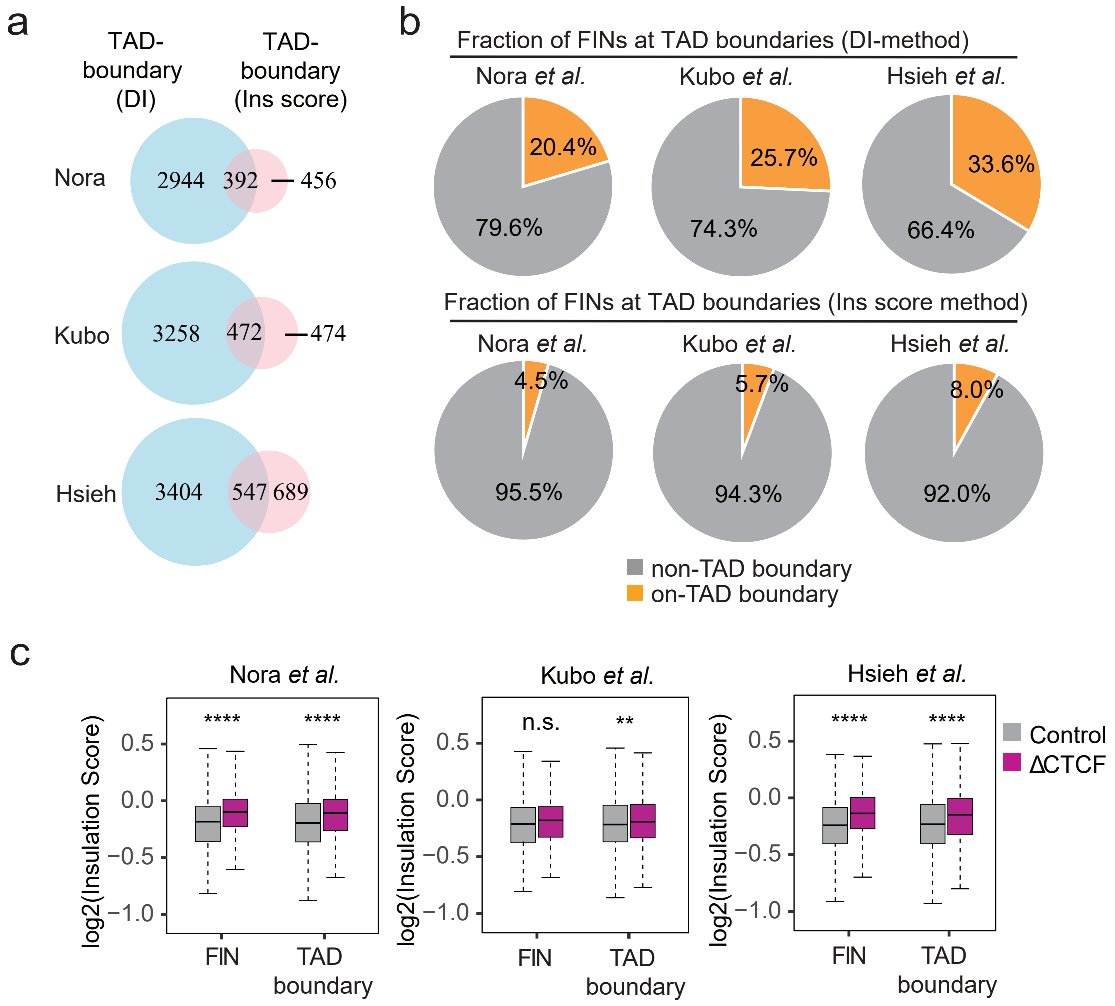

### Supplemental Figure 12

#### Supplementary Figure 12

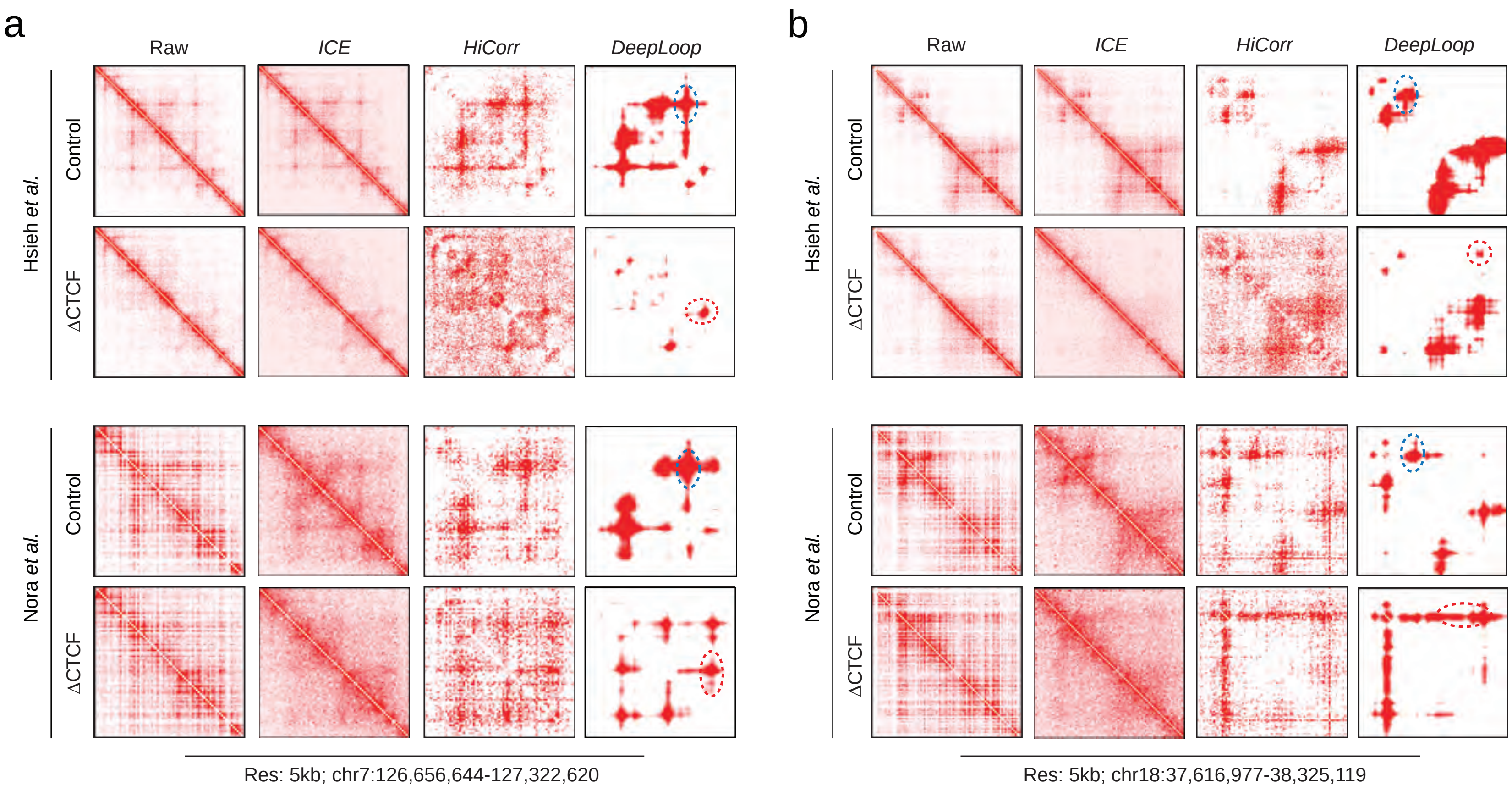
